# Synchronization of the collective air-breathing behavior in juvenile *Arapaima gigas*

**DOI:** 10.1101/2025.05.22.655545

**Authors:** Palina Bartashevich, Fritz A. Francisco, Alessandra Escurra-Alegre, Fabian Schäfer, Sven Wuertz, Jens Krause, Werner Kloas, David Bierbach

## Abstract

Animal collectives are capable of performing behaviors with high degrees of synchrony though their members might differ consistently and substantially in the focal behavior when alone. It is thus not entirely understood how these consistent differences in behavior at the individual level can be (socially) integrated into synchronized behaviors at the collective level. Here we show an unprecedented synchronized behavior in fish - the collective air-breathing of juvenile *Arapaima gigas*. Individuals of this obligate air-breathing fish from South America differed in their time between consecutive breaths when recorded alone in an aquaculture facility. However, when together in a shoal of about 200 same aged individuals, breathing is executed by a substantial portion of the shoal - within the same second. Our analysis of the individual and collective breathing patterns supported by stochastic individual-based simulations of inherently non-periodic coupled oscillators revealed that this degree of collective synchronization could be achieved by having some kind of assortative interaction rules where individuals respond towards one cluster/subgroup members stronger than to other cluster/subgroup members. By integrating this cluster synchrony rule we could successfully simulate highly synchronized collective behavior with varying proportions of otherwise diverse individuals taking part, matching our experimental observations and providing a mechanism to synchronize agents that differ consistently in the behavior in focus when in isolation.

## Introduction

Animal collectives are capable of performing behaviors with high degrees of synchrony, like instant direction changes in shoaling fish, flocking birds or walking locusts (Procaccini et al. 2011, Herbert-Read et al. 2015, Sayin et al. 2025), synchronized flashing of fireflies (e.g., (Sarfati et al. 2021)) or vocalization in frogs (Jones et al. 2014). Synchrony, e.g., the collective performance of a certain behavior at almost the same time, is often achieved when individuals follow simple social interaction rules (Couzin et al. 2002, Strandburg-Peshkin et al. 2013, Rosenthal et al. 2015) as well as possessing some degree of anticipatory abilities towards the future actions of their neighboring group mates (Couzin 2018, Bierbach et al. 2022). Interestingly, individuals in groups can synchronize behaviors for which they differ consistently among each other when observed in isolation (Jolles et al. 2020a). Causes for these consistent individual differences in behavior are manifold and comprise factors like age (and experience), sex, personality, physiology or morphology (Sih 2011, Sih et al. 2015). However, it is still not entirely understood how these consistent differences in behavior at the individual level can be (socially) integrated into synchronized behaviors at the collective level (Jolles et al. 2020a, Jolles et al. 2020b, Klamser et al. 2021), especially when it comes to trade-offs between own intrinsic behavioral rhythms or preferences and the drive to perform a certain behavior synchronously with a group in order to gain benefits (Couzin and Krause 2003, Herbert-Read 2016, McDonald et al. 2016, Jolles et al. 2017, Bevan et al. 2018, MacGregor and Ioannou 2021).

The synchronization of distinct behaviors within groups has been studied mostly in species that produce collective signals. One fascinating example is found in male fireflies that synchronize their individual light flashing to attract females (Sarfati et al. 2021, Sarfati et al. 2023). When alone, fireflies either have intrinsic regular flashing frequencies (i.e., *Pteroptyx malaccae*, see (Ramírez-Ávila et al. 2019)) or flash highly irregularly (*Photinus carolinus*). In groups, social coupling – where individuals integrate behaviors of others into their own behaviors – allows them to synchronize towards a certain group flashing frequency that can be highly periodic even when individuals alone would flash irregularly (McCrea et al. 2022, Sarfati et al. 2023). However, the synchronization of light flashing does not entail significant physiological compromise for an individual male, as it involves minimal energetic costs (Woods et al. 2007). To better understand potential trade-offs and opportunity costs of performing synchronized behaviors, it is important to study systems where individual rhythms are physiologically constrained or otherwise limited in flexibility.

In this study we investigate the air-breathing behavior in fishes that is performed both in groups and by individuals alone. Several fish species possess both an air-breathing organ as well as functional gills to take up oxygen from the water (Graham 1997, Lefevre et al. 2014, Zaccone et al. 2018). The control of the air-breathing is driven by oxygen chemoreceptors that monitor water and blood oxygen levels (Florindo et al. 2018) and thus breathing intervals of individuals of the same species are found to differ significantly among each other due to among-individual variation in metabolic oxygen demand (i.e., (Killen et al. 2018)). Although these individual differences in the underlying physiological mechanisms constrain an individual’s flexibility to adjust own breathing rhythms, several fish species exhibit synchronized collective air-breathing where individuals surface together only a few seconds apart and these breathing clusters are interspersed by minute long non-breathing periods (Kramer and Graham 1976, Chapman and Chapman 1994, Killen et al. 2018, Pineda et al. 2020). In turn, this synchrony in breathing may come at the cost of shorter breathing intervals in groups compared to when individuals are alone. For example, catfish have been shown to breathe much more frequently in groups than they would do alone, and much more frequently than would be physiologically assumed (Killen et al. 2018). These findings suggest that collective breathing may involve a degree of compromise. Given the physiological constrains of air-breathing, it is crucial to understand how individuals decide whether or not to take part in the collective breathing events, managing the underlying trade-offs. Thus, investigating possible mechanisms of collective synchrony in air-breathing fishes has the potential to inform other disciplines (e.g., behavioral ecology, neuroscience, robotics, economics) where a similar coordination of individuals that are characterized by both intra- and inter-individual differences are common scenarios.

We investigated individual and collective air-breathing of juvenile *Arapaima gigas*. This obligate air-breathing fish is assumed to represent the largest-scaled freshwater bony fish on Earth (Lüling 1964, Du et al. 2019) and inhabits wide areas of the Amazon basin (Castello 2008, Torati et al. 2019). In this more than 23 million years old species, adults are reported to exceed 3 m in size and 250 kg of weight (de Oliveira et al. 2019, Du et al. 2019, Torati et al. 2019). Juveniles are guarded by their parents in dense shoals that swim around the heads of the adults, attracted by pheromone cues (Lüling 1964). Anecdotal reports hint towards many hundreds of juveniles performing air-breathing together. Since fish become exposed to omnipresent aerial and terrestrial predators when breaking the water surface to take a breath (Kramer and Graham 1976, Kramer et al. 1983), doing this as a group might, on the one hand, even increase the detectability of the fish. Indeed, larger groups may cause greater water ripples when breathing together thus attracting surrounding predators stronger (Pacher et al. 2024). On the other hand, performing behaviors together is assumed to provide also risk-reducing advantages (Krause and Ruxton 2002, Ward and Webster 2016), especially in bird-fish interactions (Doran et al. 2022, Bierbach et al. 2025). Interestingly, it has been proposed that the frequency at which prey animals perform any behavior that exposes them to predators would be an important indicator of the predation regime the prey is facing as well as the adopted anti-predator strategy of the prey (Bednekoff and Lima 2002). When predators appear at random (such as a hunting raptor which suddenly appears and immediately attacks) regular behavioral patterns should be adopted. In turn, variability (e.g., characterized by geometric or exponential distributions) in behavioral patterns should be favored in the case of stalking predators that might be around for long periods scanning for prey to appear (see (Scannell et al. 2001, Wilson et al. 2015, Bierbach et al. 2020)). We therefore hypothesized that the time between successive breaths in our study species should follow an exponential distribution as such a strategy would imply random fish appearances at the surface, making it impossible for predators to anticipate the next surfacing event. Since earlier work on air-breathing catfish (Killen et al. 2018) imply that fish adapt their air-breathing frequencies in dependence of social context, this also leaves the possibility that Arapaima may exhibit different breathing rhythms whether they breath alone or in a group.

Our study aimed to provide the first empirical descriptions for the degree of temporal synchronization within shoals of the Arapaima. Furthermore, we adapted a generic, stochastic computational framework based on the integrate-and-fire model of inherently non-periodic oscillators (Sarfati et al. 2023) to offer mechanistic explanations of how individuals may modulate their individual surfacing needs (driven by oxygen demands) in order to achieve a large number of shoal members taking part in the synchronous air-breathing at the empirically found breathing frequencies.

To do so, we first observed single individuals as well as a shoal of ca. 200 same-aged juvenile fish in an indoor aquaculture facility in Germany (see Methods) and addressed the following questions: (1) Do individual juvenile *Arapaima gigas* differ among each other in their spontaneous air-breathing frequency when alone as known from other species (Lefevre, Wang et al., 2014; McKenzie, Burlesson, & Randall, 1991; McKenzie et al., 2015; Smatresk, Burleson, & Azizi, 1986)? (2) How strongly synchronized are breathing bouts in large shoals on a temporal scale and how many shoal members manage to take part in such synchronized breathing events? (3) Do fish in the shoal breathe more frequently than individuals alone, as known for catfish, and do breathing intervals in single individuals and/or shoals follow an exponential distribution that may hamper predators to predict future appearances at the surface (*sensu* (Bednekoff and Lima 2002))?

Using agent-based modeling, we studied (4) how the degree of intra- and inter-individual variation in individual air-breathing frequencies may influence the collective breathing performance in terms of synchrony and frequency. As there is no suitable experimental method to selectively change the air-breathing frequencies of certain shoal members yet, we simulated *in silico* shoals that were either homogenous or heterogeneous in regard to their member’s breathing rhythms (“breathing types”) and compared their collective breathing interval distributions with those observed in the live shoal. By varying proportions of distinct breathing types within *in silico* groups we asked (5) which breathing types are most influential for the collective air-breathing behavior, e.g., are their certain ‘key stone individuals’ that may dominate the collective breathing performance (Killen et al. 2018). Lastly (6), we proposed an interaction mechanism which allows individuals to minimize compromise in breathing frequencies (lowest deviation from individual breathing frequency when alone) while maximizing number of individuals taking part in the synchronous breathing events, matching our empirical observations.

## Methods

### Study organisms and sampling

We investigated juvenile *Arapaima gigas* (age about 3 months) that were imported from Peru to Germany as fingerlings and raised under indoor Recirculating Aquaculture Systems (RAS) conditions at the Manich Food Innovations GmbH, (MFI) in Dingelstedt, Thuringia, for one month prior to our observations. Fish were fed with *Chironomid* -larvae multiple times a day and kept at 28°C water temperature under 12:12 light:dark cycle throughout. We aimed at observing breathing behavior both in isolated juveniles (individual breathing) as well as in the whole shoal of about 200 individuals (collective breathing). In general, the following sequential phases of the juvenile breathing behavior can be distinguished (Fig. 1A; see (Escurra-Alegre et al. 2024) for a more complete description of Arapaima behaviors, see also SI Videos 1-2):

**Figure 1:**
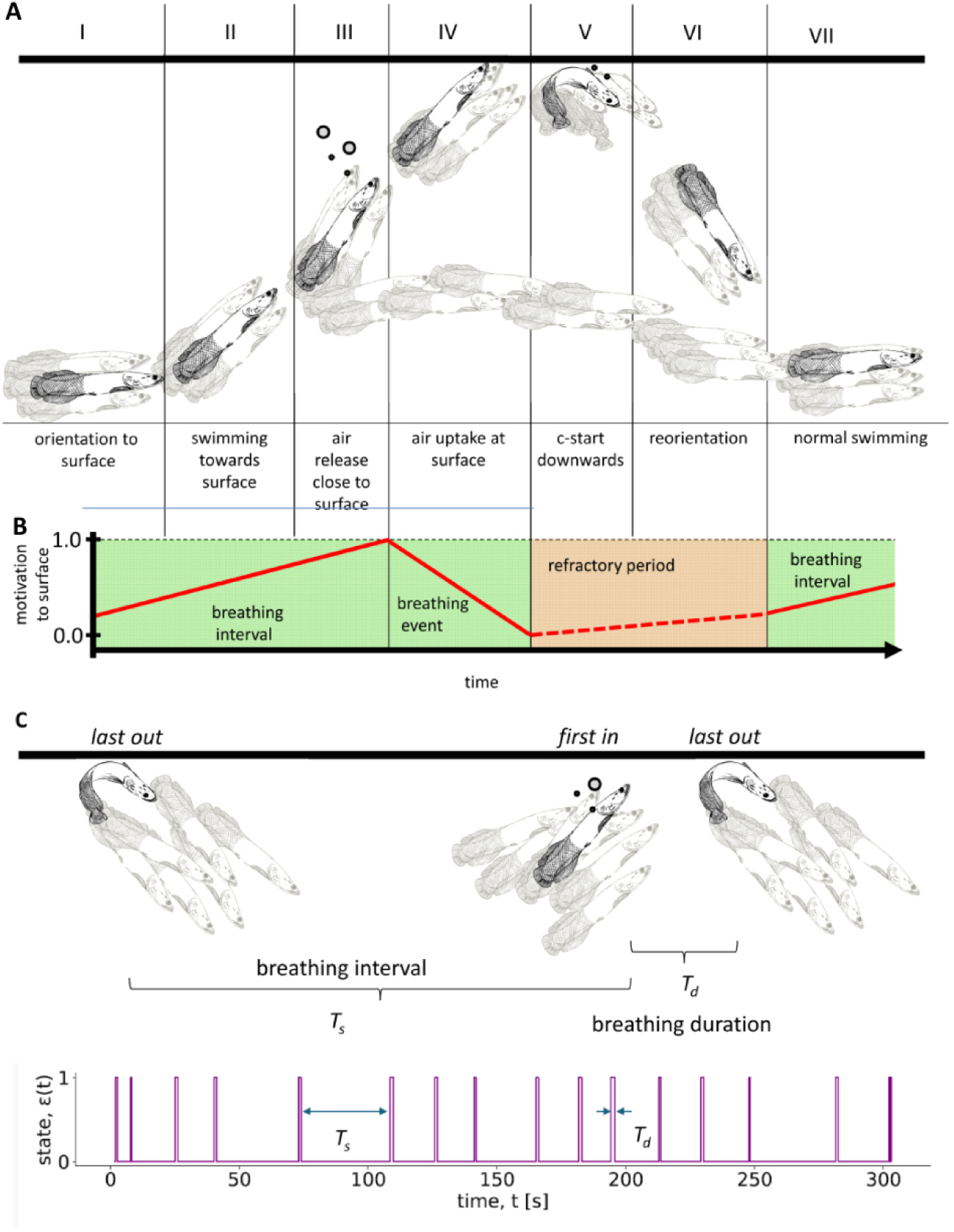
Air-breathing in juvenile Arapaima. (A) The different phases of the breathing sequence (for details, see the main text Methods). (B) Modeled motivation to surface in relation to the real breathing sequence. Note: during the refractory period fish are unresponsive to social cues (red background) thus their motivation to surface is changing only at their intrinsic rate without social influence from neighboring individuals. (c) Schematic view of the interval scoring for breathing events and the duration of the breathing. For individuals, breathing intervals T_s_ were scored as the time between an individual is diving down (phase V) and starting to release air (phase III) while breathing duration T_d_ is the time between phase III (release of air) and phase V (diving down). For the shoal, breathing intervals T_s_ were defined as the time between the last fish is diving down to the first fish is releasing air again (analogous to phase III and V), however, in the shoal different fish determined the start and end of the breathing event. Breathing duration T_d_ was scored as the time between the first fish to release air bubbles to the last fish diving down from the surface.

Phase 1 (orientation to surface): Whole shoal swims in the middle of the water column and at some point orientates towards the surface.

Phase 2 (swimming towards the surface): All fish or only fish in the upper part of the shoal swim straight or with pointed angle towards the surface. Individual fish follow each other to the surface until the first fish return form surface; when fish return into the shoal, others rarely start to move to the surface and surfacing attempts can be aborted if returning individuals are encountered.

Phase 3 (air release close to the surface): When approximately ½ to 1 body length away from surface individuals release air through gill slits well visible as bubbles. This can be considered the point of no return and we never observed any aborted surfacing attempts after an individual released air.

Phase 4 (air uptake at the surface): Individuals reach the surface and quickly swallow fresh air.

Phase 5 (c-start downwards): Immediately after the fresh air is swallowed individuals bend with a c-start like movement downwards and swim away from the surface, often with small bubbles released out of gill slits.

Phase 6 (reorientation): Individuals orientate towards others and integrate into the shoal. Phase 7 (normal swimming): The shoal swims as a whole in the middle of the water column again.

### Individual breathing

We recorded the breathing behavior of 6 individuals (body length = 29.8 cm +-0.8 cm SD) that were separated in quarantine tanks for routine check-ups at IGB Berlin. Each individual was observed twice for 1 hour for two days with one day off between observations.

Observations took place in a white, acrylic tank (100 cm x 100 cm) filled to 15 cm water level with aged tap water at 28°C. We recorded the fish with a Basler acA2040-90um camera with 2048 x 2048 pixel and 25 fps and scored the time (in frames, converted to seconds) between breathing events. We scored the breathing duration T_d_ as the time between phase III (air release close to surface) to phase V (c-start downwards). Breathing intervals T_s_ (Fig. S1) were computed as the time between an individual diving down (phase V) and starting to release the air bubbles (phase III, Fig. 1A, C).

### Collective breathing

The shoal (ca. 200 juvenile Arapaima; body size ca. 30 cm, same cohort as for individual behavior) was recorded through a glass window in their rearing tank wall (3m x 1m x 1m) with a Basler acA2040-90um, 2048 x 2048 px camera and a custom recording software (Francisco, F. A. (2025). Recording Script for USB Basler Camera (Version 1.0.0) [Computer software]. https://github.com/fritzfrancisco/ArapaimaBehaviour) at a frame rate of 25 fps (later converted to 30 fps for technical reasons). We recorded the shoal for 219 minutes in one day to avoid any major disturbances of the fish through multiple recording sessions (Fig. S2). This is important as these fish often stop feeding when subject to prolonged handling (per. observation). From the videos, we scored the duration of and the interval between collective breathing events (Fig. 1C). We defined a collective breathing duration as the time between the first fish released air bubbles on its way to the surface until the last fish was diving down from the surface. This is analogous to phase III and V for individual breathing, however, in the shoal different fish might have determined the start and end of the breathing event (first fish in - last fish out approach, see Fig. 1C). Breathing intervals for the shoal were defined as the time between the last fish diving down to the first fish releasing air again. We also estimated how many fish of the shoal took part in a breathing event. As the water turbidity did not allow counting of single individuals, we scored whether <20%, 20 - 50%, 51%-80% or >80% of the whole shoal took part. In order to establish the breathing duration of single individuals within the collective breathing events we randomly picked 30 individuals that were well visible throughout the breathing event and scored the duration of their individual air-breathing (release of old air (phase III) until diving back down from the surface (phase V)), same as was scored for individual behavior (see above).

### Statistical analyses

To estimate how much behavioral variability we encounter in the individual breathing behavior, we calculated the Repeatability of the breathing behavior that provides a measure of the among-individual variation both in breathing intervals as well as breathing durations (according to (Nakagawa and Schielzeth 2010, Dingemanse and Dochtermann 2013)). We used separate linear mixed models with trial (1^st^ or 2^nd^) as fixed factor to detect behavioral changes between the repeated test trials in intervals and durations and included individual ID as random subject factor. We used Pearson’s correlations to estimate whether breathing intervals and durations were correlated for each trial separately. Statistical tests were performed using SPSS 25 (IBM Inc.).

### The model of collective breathing

To simulate the observed collective air-breathing behavior, we use a computational model of the integrate-and-fire (IF) oscillators based on the principles of emergent periodicity proposed in (Sarfati et al. 2023). Sarfati et al. (Sarfati et al. 2023) introduced a generic, stochastic theoretical framework to explain how inherently non-oscillating individuals (i.e., with irregular periods of activity) can achieve synchronized periodic oscillations in a group.

They verified the model using the empirical data from *P. carolinus* fireflies, demonstrating that individuals converge to a common interval between flashing bursts as the group size increases. It was assumed that all fireflies in isolation have the same distribution of inter-burst intervals. However, their analytical theory shows that emergent periodicity and synchrony are guaranteed for any input single individual distribution shape of intervals between the focus events. This makes this framework well-suited for our study to model the synchronous surfacing behavior in a group of individuals with assumed consistent inter- and intra-individual variability of breathing patterns (see results too).

Following the framework of Sarfati et al. (Sarfati et al. 2023), we simulated a group of *N* individual fish whose breathing dynamics are governed by the following Equation (1):

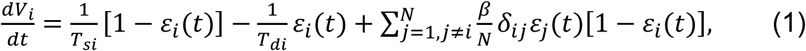

where variables V_i_ and *ε*_*i*_ characterize the internal state of an individual fish *i* = {1, . . . *N*}. The variable *V*_*i*_ represents an individual’s drive to come to the water surface to take a breath (‘motivation to surface’, see Fig. 1B), by analogy with the voltage or membrane potential in the IF of a biological neuron. It is an internal state that accumulates linearly over time until it reaches a threshold, *V*_*th*_ = 1, triggering the action, i.e., taking a breath (‘breathing event’, Fig. 1B). In other words, when an individual is not breathing, it “charges” its potential *V*_*i*_ to take a breath. This can be seen as an analogy to the decrease in blood oxygen levels that have been found to trigger air-breathing in fishes (Florindo et al. 2018). The values *ε*_*i*_(*t*) describe the binary state of the fish i at a time step t, i.e., *at the surface breathing* (discharging), *ε*_*i*_(*t*) = 1, or *underwater not breathing* (charging), *ε*_*i*_(*t*) = 0. The state *ε*_*i*_(*t*) is set to 1 as soon as the individual is fully charged (V_i_ = 1) and switches back to *ε*_*i*_(*t*) = 0 after completing the breath, i.e., when discharged to *V*_*i*_ = 0. The time *T*_*di*_ is the breathing duration of an individual i and the time *T*_*_si_*_ is the breathing interval between its two successive breathing events (Fig. 1C).

To explore how inter-individual differences influence the collective dynamics of air-breathing behavior, we consider each of the six experimentally derived distributions of individual air-breathing intervals obtained in isolation as a distinct breathing type (Fig. 2A). To simulate homogenous groups, we assigned the same breathing type to all individuals i, with T_si_values sampled from a single distribution. This will result in groups where individuals have the same breathing type. For heterogeneous groups, we considered that agents belong to different breathing types, such that we sampled *T*_*s_i_*_ values for each individual i from their own corresponding distributions. For the breathing duration, *T*_*d_i_*_, we sampled values directly from the experimental individual in the shoal breath duration distribution (see Fig. 3B). This way, both T_si_ and T_di_ are stochastic variables resampled for each agent i from the respective experimental distributions each time t the agent changes its state *ε*_*i*_(*t*). If an agent i is not breathing (*ε*_*i*_(*t*) = 0) and detects other agents j at the water surface (i.e., breathing, *ε*_*j*_(*t*) = 1), its “charging” process is accelerated by a coupling strength of *β*⁄N_*β*_, in proportion to the number of breathing fish *N*_*β*_ at a time *t* (Fig. 1B). As such, an agent *i* will reach the threshold *V*_*th*_ faster, shortening its personal breathing time *T*_*s_i_*_ compared to when being alone. We considered global network connectivity (all-to-all) between the individuals, i.e., *δ*_*ij*_ = 1 for any pair of individuals (*i*, *j*), meaning that any agent can influence and be influenced by the other.

**Figure 2:**
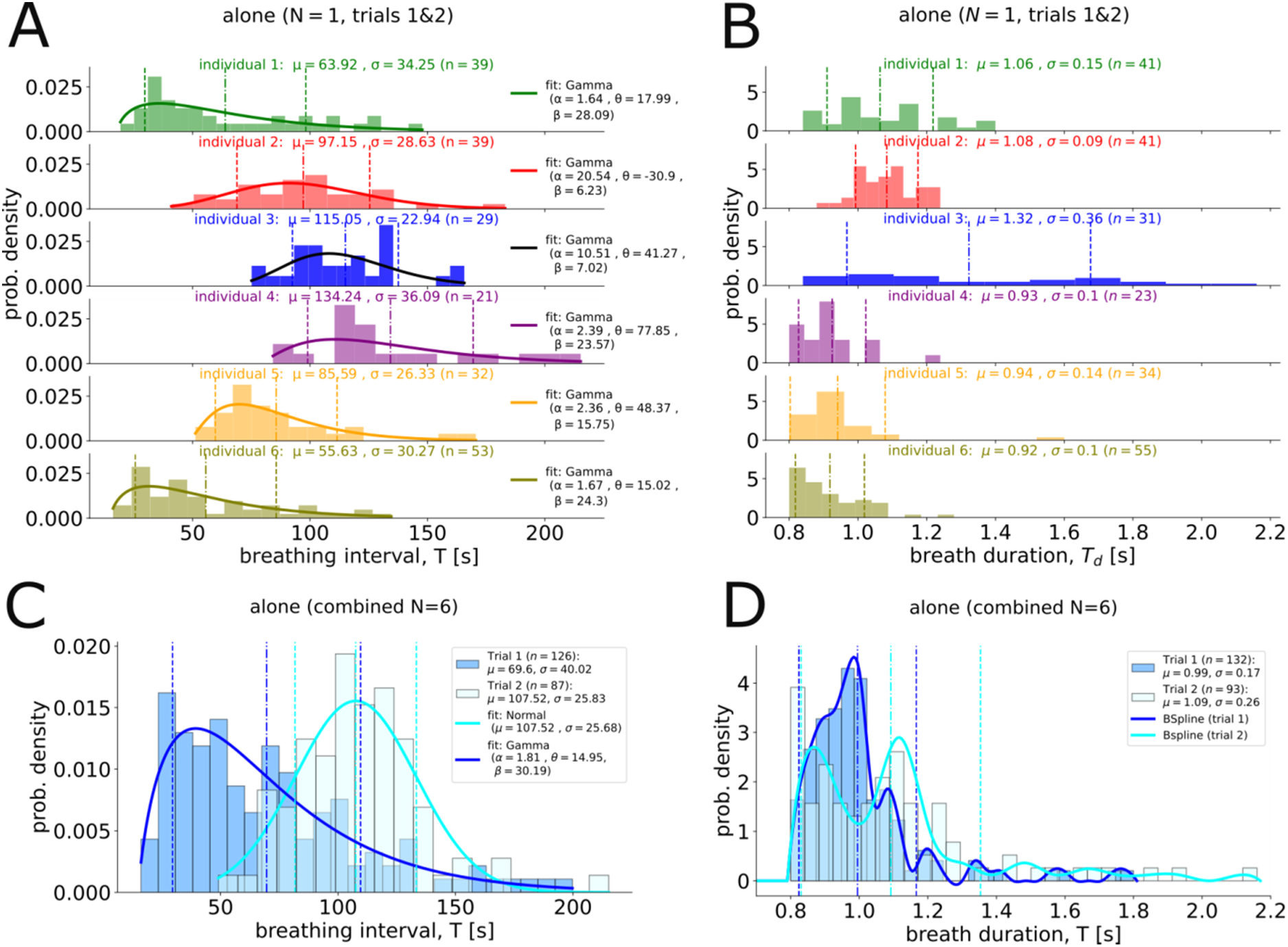
Behavioral variation in individual breathing behavior of juvenile Arapaima. Breathing intervals (A) and respective breathing durations (B) for each of the 6 individual fish recorded in isolation. Shown is the recorded data for both trials, as well as fitted Gamma distributions (KS-test always not. sig.) with shape parameters *α* > 1 (indicating that the distribution has a peak and is unimodal), location parameters *θ* and scale parameters *β* (determining the dispersion of the distribution) for each individual fish separately. The combined breathing intervals (C) and breathing durations (D) of all six individual fish recorded in isolation during the first trial (in blue) and the second trial (in cyan). The distribution of the breathing intervals during the first trial is fitted with Gamma distribution (*α* = 1.81, *θ* = 14.95, *β* = 30.19; goodness-of-the-fit: KS-statistic = 0.045, p-value = 0.95), while the second trial is fitted with Normal distribution (*µ* = 107.52, *σ* = 25.68; goodness-of-the-fit: KS-statistic = 0.079, p-value = 0.623). Solid lines in (D) show smoothed distributions of the breathing durations during the first (in blue) and the second (in cyan) trials using interpolated β-splines of degree 3 between histogram bin centers. The distribution “envelopes” formed by B-splines are normalised so that the area underneath each “envelope” sums to 1.

**Figure 3:**
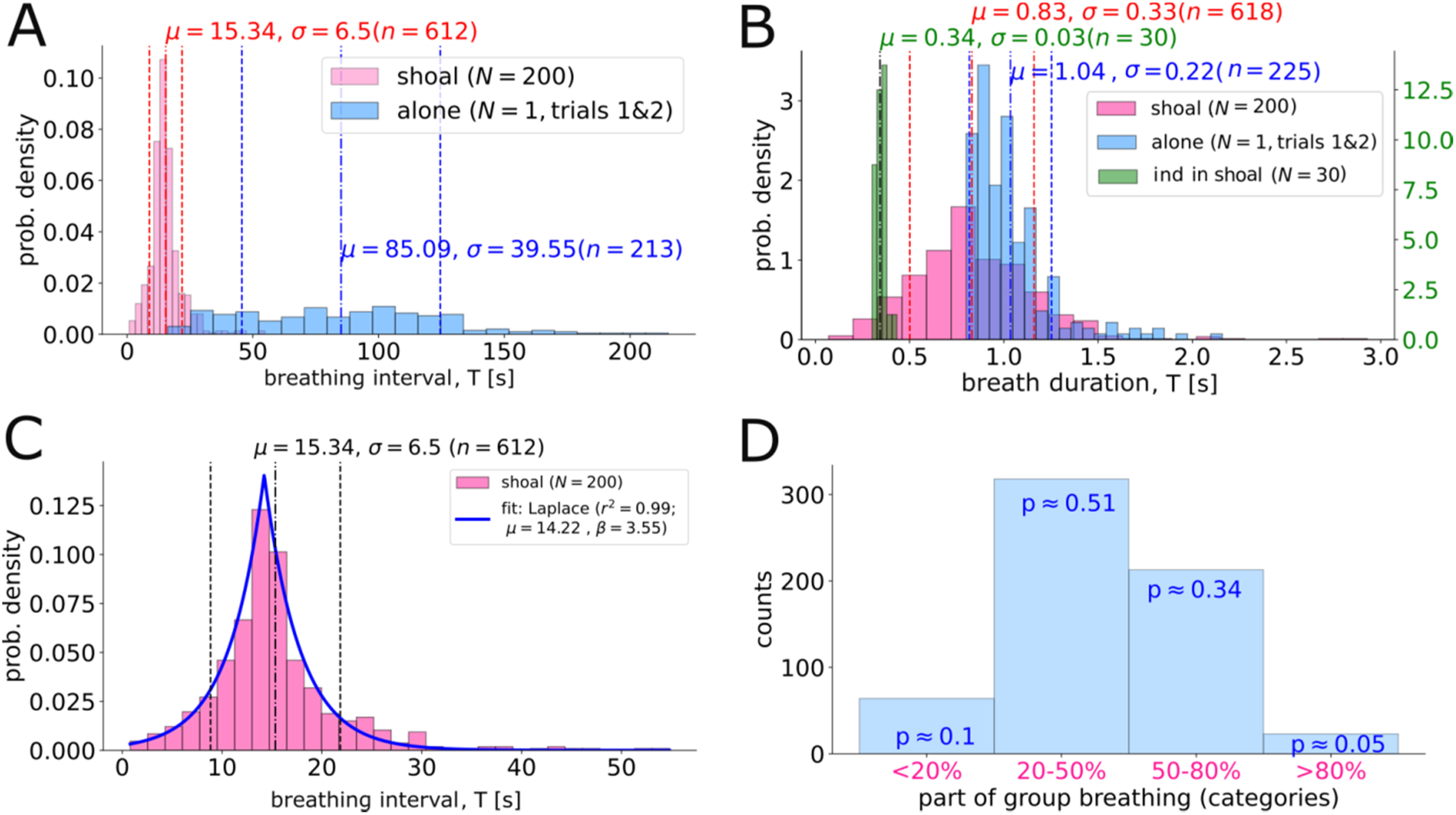
Individual and collective breathing behavior of juvenile *Arapaima gigas*. (A) Histograms of the intervals between breathing events of the shoal (in pink) and single individuals recorded alone (in blue). (B) Histograms of the breathing duration of the breathing events for the shoal (in pink), for individuals in the shoal (in green), and individuals recorded alone (in blue, both trials combined). (C) Laplace fit for the breathing intervals in the shoal with estimates for location parameter, and scale parameter as well as goodness-of-fit. (D) Proportion of the fish participating in the breathing events. Shown categories are based on the estimates that range from less than 20% fish partaking to more than 80% partaking (“<20%”, “20-50%”, “50-80%”, “>80%”).

Unlike Sarfati et al. (Sarfati et al. 2023), we introduced a refractory period, during which an individual i after finishing its breathing time *T*_*di*_ (i.e., reaching *V*_*i*_ = 0) remains socially unresponsive for the time *τ*_*ref*_. The refractory period is attributed to the stages where fish dive down from the surface and reorient into the shoal (see Fig. 1A, stages V-VI). Our exploration of the model revealed that a refractory period is required to generate the high degree of synchronization in a collective of stochastic agents employing our empirical distributions within the framework (see Fig. S5, Supplementary Note 3). To reach synchronization features similar to those of our experimental data (i.e., distribution of collective breathing intervals and shoal partaking), we used a fixed refractory period of either 1.1 s or 2.1 s, depending on the type of the model (for details see Supplementary Note 3; Figs. S5, S8).

We evaluated the performance of numerical simulations by statistically comparing the breathing interval distributions generated by the model (for different coupling strength values *β*) to the one experimentally obtained for the shoal (i.e., Laplace distribution, see Fig. 3C) using two-sided Kolmogorov-Smirnov tests. Particularly, the null hypothesis states that the simulation-generated samples follow the Laplace distribution with the same scale parameter as the experimental data (i.e., *b* = 3.55). We do not fix the location parameter *µ* in the Laplace distribution, since according to Sarfati et al. (Sarfati et al. 2023) the peak of the collective breathing interval distribution is expected to be determined by the shortest time between breathing events for an individual fish in isolation. Since the six individual distributions we use for the simulations represent a sub-sample of the entire shoal, we could have potentially missed the breathing types with such super-fast intervals, increasing the discrepancy in the comparison between distributions. To address this, we also test whether our model of synchronous breathing can reproduce a Laplace-like distribution in general, without imposing fixed parameters for the distribution.

When an individual synchronizes with others who have very different breathing patterns compared to its own, it has to adjust its individual natural rhythm stronger than when synchronizing with those of a similar breathing type. We assume this mechanism would then promote spatial self-sorting within the shoal based on breathing type (forming ‘breathing clusters of similar breathing types’, see for example (Croft et al. 2009, Jones et al. 2010, Szorkovszky et al. 2018)). Due to self-sorting, spatial constraints would limit immediate interactions between individuals from different breathing clusters. Under these conditions, we expect to get partial synchronization (where only parts of the shoal participate in a synchronized breathing event) with *some degree* of compromise to emerge, which by definition will be less than in the case of full synchronization (where the whole shoal takes part in a breathing event = non-clustered heterogeneous shoals). To quantify how well the simulated partaking rates based on the above mechanism match our empirical observations of partaking rates, we computed the sum of squared differences (SSD) between the empirical probabilities of occurrence of each partaking category (i.e., “<20%,” “20-50%,” “50-80%,” “>80%”; see Fig. 3D), which represent the proportion of the shoal surfacing, and the corresponding probabilities from the simulation. Furthermore, we analyzed the level of individual compromise for simulated homogenous, heterogenous with varying degree of clustering, and non-clustered heterogeneous groups in relation to the fraction of simultaneously breathing individuals (Fig. 7). Individual compromise is calculated for each breathing event as the difference between an individual’s simulated breathing interval in a group and the individual’s empirical average breathing interval when alone (in seconds).

## Results

### Variability in individual breathing behavior of juvenile *Arapaima gigas*

Individuals differed consistently among each other in the intervals between spontaneous air-breathing when alone (sig. Repeatability of 0.53 [0.27-0.77], LRT: *chi²*=117, *P*<0.001, Fig. 2A). We further found a significant habituation effect with longer breathing intervals during the second trial (avg. first observation: 69.6 s ± 40.0 SD; avg. second: 107.52 s ± 25.83; sig. effect of trial in LMM: *F*_1,206_=99.8, *P*<0.001; Fig. 2C). Similarly, the duration of the air-breathing behavior was different among individuals (sig. Repeatability of 0.45 [0.21 −0.71], LRT: *chi²*=89, *P*=0.01, Fig. 2B) and fish took on average longer to breathe in the second trial (avg. first trial: 0.99 s ± 0.17 SD; avg. second trial: 1.09 s ± 0.26; sig. effect of trial in LMM: *F*_1,218_=16.9, *P*<0.001, Fig. 2D). Although we found evidence for the presence of significant individual differences in breathing intervals, we can further classify the individuals as either fast (id 1 and 6), medium (id 2 and 5) or slow (id 3 and 4) breathers (referred to as “breathing types”) based on their average breathing intervals (Fig. 2A). For both trials, air-breathing durations and subsequent intervals were positively correlated (Pearson’s *r*; first observation: *N*= 126, *r*=0.282, *P*<0.001, Fig. S3A; second observation: *N*=87, *r*=0.276, *P*=0.01, Fig. S3B), indicating that on average a longer time since the last breath led to a longer subsequent breathing duration.

### Synchrony in collective breathing behavior of juvenile *Arapaima gigas*

In the observed shoal, we found individual fish to perform very short air-breathing events; three times shorter than the air-breathing duration of the individually observed fish (individual in shoal: 0.34 s ± 0.03 SD, N= 30; individual alone: 1.03 s ± 0.22, Fig. 3B). We recorded 618 collective breathing events and in more than 85% of all events a fraction of 20% to 80% of the fish took part in the collective breathing (Fig. 3D). Individuals were swimming most of the time very close in one shoal in a wall or cube like formation (SI video 1) and fish closest to the surface were taking part in the air-breathing (Fig. 1A; SI video 1 & SI video 2). The duration of a collective breathing was on average 0.83s (±0.33 SD) long (Fig. 3B) and the more fish taking part the longer the bout (Spearman’s *r*_duration-partaking_ = 0.302, *p*<0.001, *N*=618; Fig. S4A). This shows that even when about 100 individuals taking a breath together (=50% of the shoal), this just lasted for about as long as it would take a single individual in isolation to take a breath alone (Fig. 3B). This further exemplifies the extreme degree of synchronization of the collective air-breathing in this species. Intervals between collective breathing events were on average 15.34 s (± 6.5s SD), thus much shorter as found for individuals alone (Fig. 3A). There is also a positive correlation between the proportion of the shoal taking part in a breathing event and the length of the subsequent collective breathing interval (Spearman’s *r*_interval-partaking_ = 0.09, *p*=0.021, *N*=612, Fig. S4B), indicating that the more fish took part the longer the shoal waits until the next breathing is initiated. The distribution analysis of the individual and collective breathing intervals revealed that individual breathing follows Gamma distributions (Fig. 2A) while the collective breathing shows strong evidence for a Laplace distribution (i.e., double-exponential distribution, Fig. 3C, *r*² =0.99).

### Agent-based simulations of air-breathing (surfacing) coordination in groups

First, we considered a group of N=200 agents where all individuals shared the same breathing type (homogeneous group). Here, we sampled for each agent i individual breathing intervals *T*_*si*_ from one of the Gamma distributions derived from the experimental data of isolated individuals (Fig. 2A), corresponding to the given breathing type. Depending on the breathing type, individuals exhibited different breathing interval characteristics within the group (Fig. 4B-E). As the strength of social interactions (coupling strength *β*) increased from 0 to 1, individuals started to synchronize their breathing (see Fig. 4A, Fig. S5) such that the individual breathing intervals initially increased (Fig. 4B) while their standard deviation showed a sharp decrease (Fig. 4C). With further increase of coupling strength *β* > 1, this trend was reversed as the mean breathing interval decreased (Fig. 4B) and its standard deviation increased (Fig. 4C), regardless of the breathing type. This means that with a strong coupling strength among individuals the synchronous collective breathing pattern became more irregular rather than periodic, i.e., it did not follow a fixed interval (e.g., see Fig. 4A).

**Figure 4:**
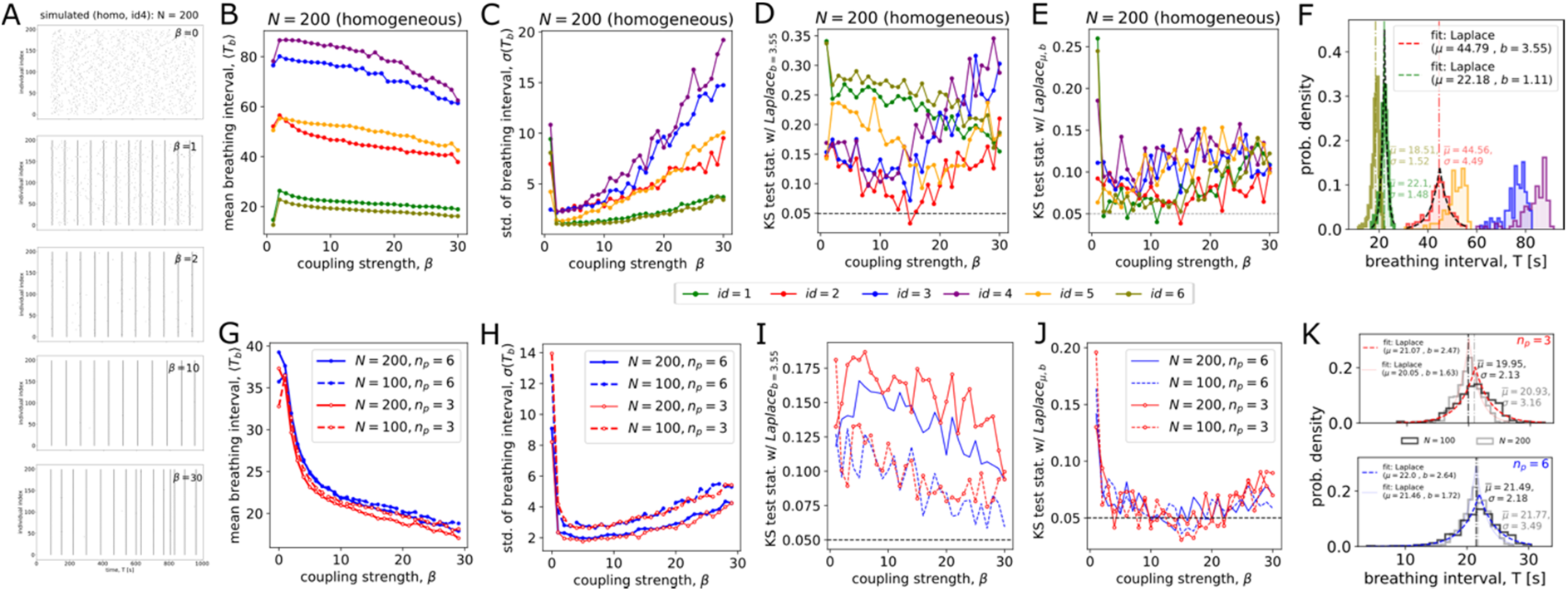
Effect of individual breathing diversity on the collective air-breathing in simulated homogeneous and heterogeneous groups. (A) Example of simulated breathing intervals for a group (N=200) composed of only id=4 breathing types at different social coupling strengths *β*. (B) Simulated mean breathing intervals and (C) their standard deviation for homogenous groups composed of either one of the 6 individual breathing types. (D) Two-sided Kolmogorov-Smirnov test results verify at each β whether breathing intervals in the modelled groups come from the Laplace distribution with the empirical scale parameter *b* = 3.55 or (E) from any Laplace-like distribution without fixed parameters. The KS-test values above 0.05 indicate rejection of the null hypothesis, meaning the collective breathing intervals of the simulated group do not follow a Laplace distribution. (F) Depiction of the model-generated collective breathing interval distributions for six homogenous independent groups with β parameters corresponding to their best fits to the Laplace distribution (*β* = 11 for id ∈ (1,3,4,6) and *β* = 15 for id ∈ (2,5)). (G) Simulated mean breathing intervals and (H) their standard deviation for heterogeneous groups composed of either only *n*_*p*_ = 3 or all *n*_*p*_ = 6different individual breathing types for group sizes of *N* = 100 and *N* = 200. (I) KS test results verify at each β whether breathing intervals in the modelled groups come from the Laplace distribution with the empirical scale parameter *b* = 3.55 or (J) from any Laplace-like distribution without fixed parameters. (K) Depiction of the model-generated collective breathing interval distributions for heterogeneous independent groups of *n*_*p*_ = 3and *n*_*p*_ = 6with *β* = 14 for group size of *N* = 100; as well as for group size of *N* = 200 (here with *β* = 15 in case of *n*_*p*_ = 3and *β* = 13 in case of *n*_*p*_ = 6). All distributions were generated from 10 simulations, 1000 time steps each with *dt* = 0.01 and the refractory period *τ*_*ref*_ = 1.1.

In the next step, we compared the breathing interval distributions generated by the model for different coupling strength *β*with the experimentally obtained data for the shoal (see Methods for details). While we found some breathing types in a homogeneous group to follow Laplace-like distributions in their collective breathing intervals, either the location parameter (breathing type id=2: *µ* = 44.79, Fig. 4D, F) or the scale parameter (breathing type id=1: *b* = 1.11, Fig. 4E, F) did not match with empirical data (see Supplementary Note 1). Notably, the Laplace-like distribution of breathing intervals was observed only for strong social coupling between individuals (*β* = 11 for breathing type id=1 and *β* = 15 for breathing type id=2).

Since simulations of groups with homogeneous breathing types did not match with the empirical collective breathing interval distribution, we simulated heterogeneous group compositions where all six given breathing types were included. We also tested whether the three main breathing types (e.g., id=6 as fast; id=5 as medium; id=4 as slow; for details see Supplementary Note 2: semi-heterogeneous case) instead of all six can be sufficient to represent the empirically found collective breathing intervals. By studying the statistical properties of simulated groups of N=100 and N=200 individuals – each composed of either all six or only three breathing types, with each type in equal proportion relative to a group size – our simulations showed that as social coupling strength increased the mean breathing interval in heterogeneous groups declined more rapidly compared to the homogeneous groups (Fig. 4G vs Fig. 4B). This can be explained by a convergence of the breathing interval to the fastest breathing type present within the group as exemplified by our analysis of semi-heterogeneous groups (i.e., groups containing members with just two distinct breathing types; see Supplementary Note 2: semi-heterogeneous case). The standard deviation of the breathing interval in the group with diverse (i.e., more than 2) breathing types tended to increase slower with increasing coupling strength compared to the homogenous groups (Fig. 4H vs. Fig. 4C) while it in general followed the pattern of the fastest breathing type in the group (i.e., id=6 in Fig. 4C; also see Supplementary Note 2: semi-heterogenous case).

We found collective breathing dynamics differ between group sizes but not between six- and three-breathing type compositions (Figs. 4G-J), most probably because within the underlying 6 individual breathing types there were already 3 distinct main breathing types with 2 individuals each (see also semi-heterogeneous cases in Supplementary Note 2). Consistent with (Sarfati et al. 2023), the mean and the standard deviation of the group breathing intervals increased with decreasing group size (Fig. 4G, H).

All considered heterogeneous group compositions generated Laplace-like distributions of collective breathing intervals (Fig. 4J). However, none of them matched the distribution of breathing intervals with empirical scale parameter *b*, regardless of the coupling strength *β* (Fig. 4I), and characterized by narrower distribution shape overall (Fig. 4K). This indicates that the included breathing types may not represent the full range of breathing types present in the observed Arapaima shoal. Notably, our selected breathing types for the three-types groups did not follow any Laplace-like distribution in a homogenous case (i.e., id ∈ (4, 5, 6), also see Fig. 4E) but did so when analyzed as heterogeneous group (Fig. 4J). Despite this, when interacting together, the breathing dynamics of such three-breathing types compositions resembled those of the six-personality composition (*n*_*p*_ = 3 vs. *n*_*p*_ = 6 in Fig. 4K), which includes breathing types that can exhibit a Laplace-like distribution in the homogenous case on their own (i.e., due to the intra-individual variation). This suggests that the Laplace-like distributions of the distinct behavior in focus - collective synchronous air-breathing - are an indication of a heterogeneous group member composition in regard to the individually performed behavior.

### Impact of varying breathing type ratios on the collective air-breathing in heterogeneous groups

Above, we demonstrated how individual differences – both within and between individuals – affect collective air-breathing behavior, assuming equal proportions of each breathing type within the group. Here, we explored how varying these proportions will affect the collective air-breathing dynamics by adjusting the extremes – decreasing the number of “slow breathers” and increasing the number of “fast breathers”. We focused on the three-breathing type group compositions, sampling individual breathing intervals from the fitted Gamma distributions of individuals 4, 5, and 6 when tested alone (see Fig. 2A). We maintained the proportion of “medium” (id=5) breathers at ⅓ of the group size *N* and varied the number of “’fast” (id=6) breathers *N*_*fast*_ from 0 to ⅔. The number of “slow” (id=4) breathers is respectively adjusted to be the remaining fraction (⅔\**N* − *N*_*fast*_).

The mean breathing interval of individuals in a group decreased asymptomatically as the number of fast breathers was increased (Fig. 5A). The standard deviation of the breathing intervals initially increased with the introduction of fast breathers, peaking at 10 individuals in case of N=200 and at 3 individuals for N=100. As more fast breathers were added, the standard deviation started to decrease, indicating that the group breathing became more regular with addition of fast breathers (Fig. 5B). Once there are 10 fast breathers in a group of either N=100 or N=200 individuals, their influence on the variability of the breathing intervals became comparable between the considered group sizes (Fig. 5B).

**Figure 5:**
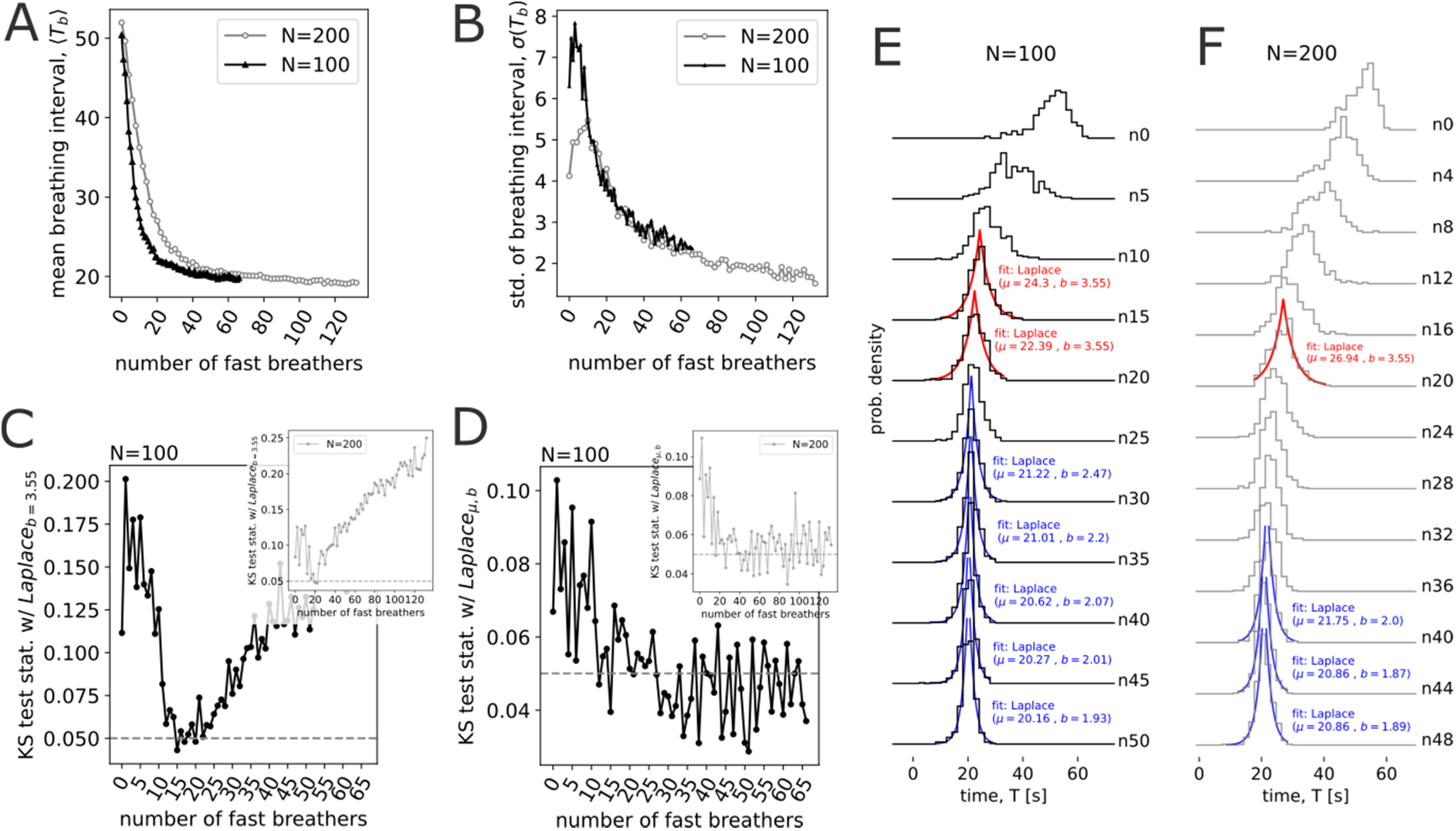
Impact of varying ratios of breathing types on the collective air-breathing in simulated heterogenous groups. (A) Simulated individual breathing intervals (means) and (B) standard deviations in a heterogeneous three-breathing-type group composition as a function of the number of “fast” breathers *N*_*fast*_, for different group sizes N. The number of present individuals of each of the types is re-calculated as (*N*_*fast*_, ⅓\**N*, ⅔\**N* − *N*_*fast*_) for “fast”, “medium”, and “slow” breathing types, respectively. (C) Matching of simulated distributions of collective air-breathing intervals to the empirically found Laplace distribution with the scale parameter *b* = 3.55 or (D) to any Laplace-like distribution without fixed scale parameter. (E, F) Examples of the simulated breathing interval distributions for groups of N=100 (E) and N=200 (F) individuals, with increasing numbers of “fast” breathers (e.g., n0 corresponds to their absence). In (A)-(F), the strength of the social coupling is set to *β* = 14. All distributions were generated from 10 simulations, 1000 time steps each with *dt* = 0.01 and the refractory period *τ*_*ref*_ = 1.1.

We found that when proportions of fast breathers started to exceed about 10% in a group of *N* = 200 and 15–20% in a group of *N* = 100 the collective breathing intervals matched Laplace-like distributions (Figs. 5D-F). However distributions with the empirical scale parameter *b* = 3.55 were only found for groups with 10%-15% fast breathers (Figs. 5C, E-F). Notably, when there are no “fast” breathers in a group and “slow” breathers are in the majority (i.e., ⅔ of N), the group with “medium” breathers (that are now the fastest breathers in the group) did not follow a Laplace-like distribution of breathing intervals (Fig. 5D). This means, that a small minority of “fast” breathers – around 10% to 15%, depending on group size – not only replicated the shape of the empirical Laplace distribution for breathing intervals but was also sufficient to significantly accelerate the collective breathing rate, bringing it closer to the empirical location parameter (i.e., *µ* = 14.22; Fig. 3C). However, the remaining deviation from the empirical mean (i.e., *µ* = 26.94 in the case of *N*_*fast*_ = 20, *N* = 200; Fig. 5F) implies that the “fast” breathing type we consider in our simulations might not represent the absolute “fastest” individual type present in the empirical shoal.

### Individual compromise and collective partaking rates

While our simulations showed how the empirical Laplace distribution of the collective breathing intervals can emerge in a shoal with heterogeneous individual breathing types, this scenario assumes that all individuals participate in the breathing events (100% partaking). In contrast, our observations of the live fish showed that typically only a fraction of the individuals and not the entire group came to the surface (partial partaking) (Fig. 3D, SI video 1 & 2). This fraction - in the majority of cases between 20% to 80% of the group - form a synchronized subgroup (cluster), while the fraction of those that do not participated in the surfacing can be seen as a separate subgroup. This behavioral pattern can be referred to as *cluster synchronization*, where the group splits into distinct clusters that exhibit internal synchrony but are not synchronized with other clusters (e.g., (Barreto et al. 2008, Kato and Ishii 2024, Lodi et al. 2024)). To simulate this, we assume a fully connected network where individuals of each breathing type (fast, medium, slow) form a breathing cluster and the strength of interactions is weaker between individuals of different breathing clusters (inter-cluster coupling, *β*_*inter*_*clu*__) and significantly stronger within the same cluster (intra-cluster coupling, *β*_*intra*_*clu*__; such that *β*_*intra*_*clu*__ ≫ *β*_*inter*_*clu*__).

We identified a transition region at *β*_*inter*_*clu*__ = 0.7 where the collective breathing behavior dramatically changes. Below the transition (*β*_*inter*_*clu*__ ≪ 0.7), individuals primarily breathe together with individuals of their own breathing types, making the “20-50%” partaking category the most dominant (i.e., with 0.88 probability of occurrence, see Fig. 6D). Since collective breathing intervals are measured from the last individual to dive in one cluster to the first one that is resurfacing (from any cluster), this leads to short but highly inconsistent intervals between breathing events, and thus high standard deviation of the breathing intervals (“D” in Fig. 6C). As a result, the collective breathing intervals failed to form a Laplace-like distribution, regardless of the coupling strength *β*_*intra*_*clu*__ (Fig. 6B). Above the transition (*β*_*inter*_*clu*__ ≫ 0.7), our simulations showed a strong tendency for full synchrony of all individuals in the group during the breathing events (indicated by the increasing dominance of “>80%” partaking category; Fig. 6F), especially when the strength of the inter-cluster coupling exceeded *β*_*inter*_*clu*__ = 1.0. The standard deviation of the breathing intervals and the shape of their Laplace-like distributions did not change with increasing inter-cluster coupling strength after the transition (Fig. 6F, G). Most notably our empirical data is best matched around the transition region. Here, the lowest SSD values are found which indicates the emergence of cluster synchronization with proportions of partaking individuals comparable to the empirical estimations (“E” and “G” in Fig. 6A). Also, Laplace-like distributions of breathing intervals emerge above the transition region (i.e., for *β*_*inter*_*clu*__ > 0.6 in Fig. 6B). In summary, at weak inter-cluster and strong intra-cluster coupling (i.e., *β*_*inter*_*clu*__ = 0.7, *β*_*intra*_*clu*__ = 15) our model showed empirical-like shoal part-taking behavior in the breathing events with a Laplace-like distribution of collective breathing intervals. Nevertheless, regardless of whether the group exhibited full or clustered partaking, the simulated distributions of collective air-breathing intervals consistently differ from the empirical data, showing either narrower or shifted Laplace-like distributions compared to the empirical fit (Fig. 6G). This suggests that the diversity in the shoal may be higher than three breathing types inferred from isolated experiments, which were used in the simulations (see also above).

**Figure 6:**
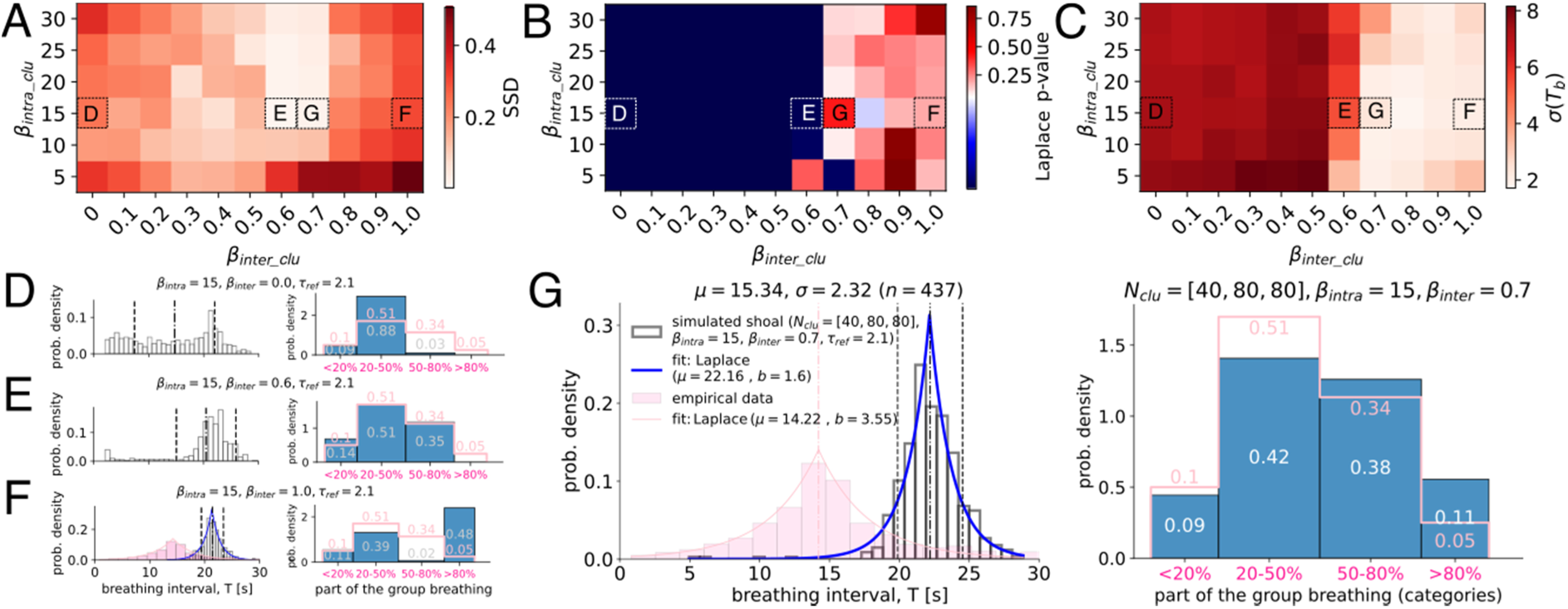
Influence of intra- and inter-cluster coupling strength on the proportion of part-taking individuals and breathing interval distribution in simulated heterogeneous groups. (A) Difference between empirical and simulation-generated probabilities of the proportions of individuals participating in breathing events depicted as the sum of squared differences (SSD) for varying intra- and inter-cluster coupling strengths. (B) Likelihood of encountering a Laplace-like distribution for different combinations of the intra- and inter-cluster coupling strengths based on the KS test statistics. If the p-value is below 0.05 (in blue), the intervals do not follow a Laplace-like distribution. (C) The standard deviation of the simulation-generated collective breathing intervals for varying intra- and inter-cluster coupling strengths, indicating the degree of irregularity in the breathing pattern. (D-G) Histograms of the distributions of collective breathing intervals and proportions of the group participating in the breathing event in simulation (in blue) versus empirical data (in pink) for four different sets of parameters, highlighted by different corresponding letters in (A-C). All distributions were generated from 10 simulations, 1000 time steps each with *dt* = 0.01 and the refractory period *τ*_*ref*_ = 2.1. Simulated groups were composed of *N* = 200 individuals from three breathing types with ids ∈ (4,5,6) and different intra- and inter-cluster coupling strengths. Based on the results of the previous section, we assumed fast breathers are in the minority and represent the group with an unequal ratio of fast:medium:slow breathers as 1:2:2, corresponding to breathing cluster sizes of N_clu_ ∈ (40,80,80) agents.

**Figure 7:**
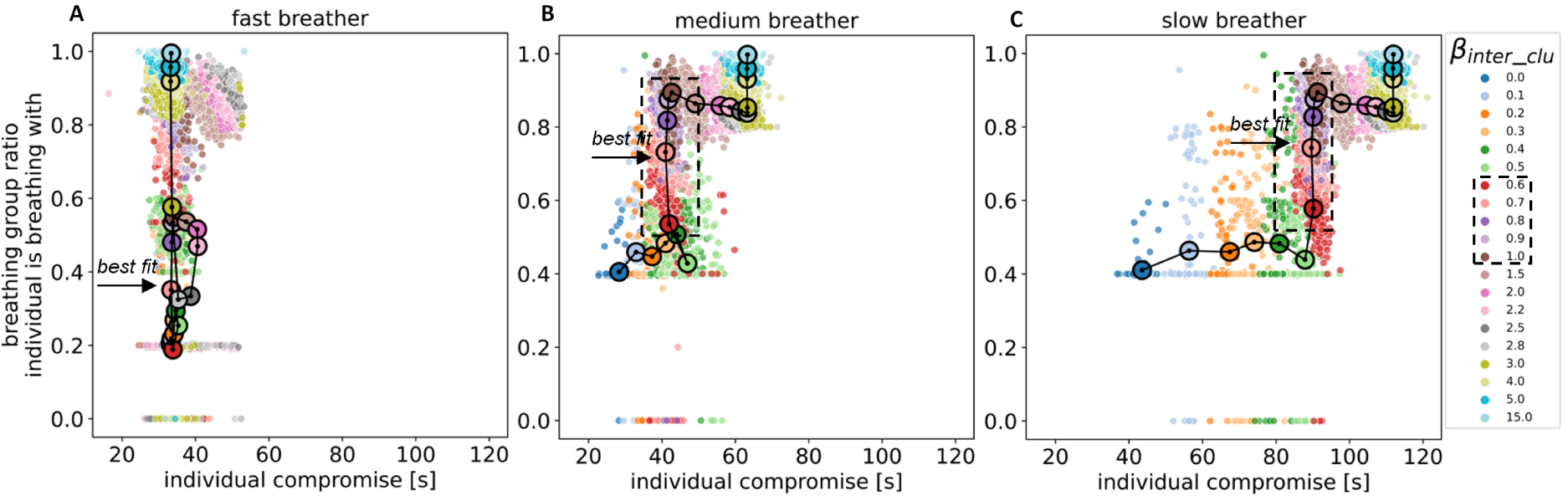
Partaking in simulated collective breathing as a function of individual compromise. Individual compromise (the deviation of an individual’s breathing interval in a group from its mean breathing interval when alone) is plotted against the number of individuals an individual is breathing synchronously with [given as ratio of the whole group]. This relationship is analyzed under different inter-cluster coupling strengths *β*_*inter*_*clu*__, while keeping high intra-coupling strength *β*_*intra*_*clu*__ = 15. In a simulated group of N=200 individuals with a 40%-40%-20% composition of slow, medium, and fast breathers (see Fig. 5), we tracked one randomly selected slow (A), one medium (B), and one fast (C) breather. At *β*_*inter*_*clu*__ = 0 individuals of each breathing type are only responsive to others of the same breathing type (independent homogeneous groups), thus partaking ratios mirror fraction of individuals of this breathing type in the shoal (40-40-20) with some breathing events that coincide with other breathing types by chance. Above *β*_*inter*_*clu*__ = 5 individuals of all breathing types breathe together (full partaking, i.e., non-clustered heterogeneous group). Big circles indicate mean values in the individual compromise and partaking rate for different *β*_*inter*_*clu*__. The arrow points to the simulations which matched best with empirical data in terms of partaking and collective breathing interval distribution.

When looking at an individual’s compromise (the deviation of an individual’s breathing interval in a group from the mean breathing interval of this individual breathing type when alone) in relation to the proportion of others that synchronize their air-breathing with this individual, our simulations show that slow breathing types can adjust the partaking rate by changing inter-cluster responsiveness *β*_*inter*_*clu*__ (Fig. 7A). This allows individuals to shift from breathing synchronously almost exclusively with others of their own breathing type at inter-cluster responsiveness *β*_*inter*_*clu*__ = 0 to breathing in full synchrony with all shoal members regardless of breathing types at high inter-cluster responsiveness values (Fig. 7A-C). At inter-cluster responsiveness around the aforementioned transition region (*β*_*inter*_*clu*__ ∈ [0.6,1]) every breathing type in the group can adjust partaking rates without additional costs/compromise (see a sharp jump highlighted by a dashed rectangle in Figs. 7A-C). This means that in this region, even minute changes in responsiveness lead to immense changes in partaking rates while individual compromise is maintained at a similar level for all simulated breathing types. Not surprisingly though, the range of possible compromise was the smallest for the fast breathing type (Fig. 7C) and the largest for the slow one (Fig. 7A).

This can be explained by the fact that fast breathers cannot extend their breathing intervals as much as slow breathers that could easily breathe at faster intervals. For fast breathers, the inter-cluster responsiveness only determined the partaking rate, while for slow and medium breathing types inter-cluster responsiveness also affected the individual compromise. Furthermore, above a certain inter-cluster responsiveness (*β*_*inter*_*clu*__ > 0.8), slow and medium breathers have not been found to breath alone while this could still happen for fast breathers (for details see SI, Table S1).

On the plots, we highlighted the parameter regime around the transition region (*β*_*inter*_*clu*__ ∈ [0.6,1]) by the dashed rectangle. The results in (A)-(C) are based on 10 simulations, each running for 1000 time steps with *dt* = 0.01 and the refractory period *τ*_*ref*_ = 2.1.

## Discussion

Our study provides unprecedented evidence for the synchronous collective air-breathing behavior of juvenile *Arapaima gigas*. In a shoal of about 200 individuals kept in an aquaculture facility we found often more than 100 individuals surfacing together within a second, in several cases also all 200 individuals. The shoal repeated this highly synchronized behavior almost every 15 seconds while individuals observed alone showed consistent differences in the frequency of the air-breathing behavior that ranged from about 1 minute to more than 2 minutes. Interestingly, when comparing the distribution of the breathing intervals between isolated individuals and the shoal, we found individual breathing intervals to follow broad gamma distributions while as a collective the intervals were Laplace distributed and thus more regular than for individuals alone. Our simulation results illustrated that a *heterogeneous* (in terms of individual breathing intervals) shoal composition is necessary to match the Laplace-like experimentally observed distribution of breathing intervals in a group of *N* = 200 fish. Especially the “fast” breathers seem to be important drivers of the observed interval distributions with 10% of fast-breathing individuals are enough to significantly accelerate the breathing rhythm of the whole group of 200 to the interval of fast breathers. Using our model, we provide further evidence that the partaking rate between 50% and 80% of the shoal members is likely due to a tendency of individuals to breathe within subgroups of similar breathing intervals due to self-sorting and spatial constraints in the shoal - a phenomenon called cluster synchronicity. As the surfacing exposes Arapaima to aerial predators, individuals when alone may breathe less often and at less predictable intervals while the more regular Laplace distributed breathing in the shoal might be caused by compromises needed to allow individuals with different individual breathing needs to still breath together as a group - a known anti-predator strategy as well.

Juvenile Arapaima in our study differed vastly in their individual breathing intervals when observed alone which points towards among-individual variation in metabolic oxygen demands, similar to what is known from catfish (Killen et al. 2018). Nevertheless, individual breathing intervals had a broad range which seemingly allowed the fish to synchronize their air-breathing when together in a shoal with often about 50% and more of the shoal members breathe at the same time. This temporal synchrony is unprecedented as other fish species do not show such high temporal coordination in air-breathing (Kramer and Graham 1976, Chapman and Chapman 1994, Killen et al. 2018). Perhaps this is due to the fact that other reports stem from non-shoaling or facultative shoaling species, while Arapaima juveniles have been found to strongly shoaling and even schooling, thus are close together when the air-breathing is initiated. As found in catfish (Killen et al. 2018), individuals in the shoal breathe much more frequently than they would do alone letting us assume a strong social coupling and making air-breathing in Arapaima - similar as proposed in catfish - an emergent collective behavior (Couzin and Krause 2003) and an example of social facilitation of behavior (Webster and Ward 2011).

Distributions of air-breathing intervals of single individuals followed a broad gamma distribution, while those exhibited in the shoal matched a broadened Laplace distribution. The intervals of a behavior that exposes prey to predators are an important indicator of both the predator’s anticipated encounter rates as well as the prey’s anti-predator strategy (Scannell et al. 2001). For juvenile Arapaima, visiting the surface exposes them to aerial predators that might always be around in their natural habitats looking for prey to appear. Thus, the observed surfacing at low frequency and at irregular and unpredictable intervals seems to be an adaptive anti-predator strategy for single individuals. For the shoal, however, the surfacing pattern is different and intervals between subsequent breathing events are more regular. Compared to exponential distributions like gamma, a Laplace distribution has a sharp peak (Abramowitz and Stegun 1964) indicating that the group tends to breathe most frequently at a certain interval, what makes their surfacing behavior potentially more predictable to predators. However, in contrast to fireflies, where their collective flashing might serve as a signal and is thus believed to be most effective if regularly and periodically (Smith and Harper 2003, Bro-Jørgensen 2010), the collective execution of the air-breathing has most likely an anti-predator function and we thus hypothesize that the anti-predator benefits of performing the surfacing as a large group outweighs the costs of a higher and more regular breathing frequency. A surfacing shoal might be well visible for any predator that is present anyway (Doran et al. 2022) - regardless of the frequency range and regularity of the breathing intervals but predators maybe impeded due to confusion or deterrence effects that come along with collectively performed behaviors (see (Landeau and Terborgh 1986, Magurran 1990, Bode et al. 2010, Doran et al. 2022, Bierbach et al. 2025).

In order to explore possible mechanisms of how Arapaima shoals are able to synchronize the majority of individuals that themselves differ vastly in breathing intervals to perform the air-breathing collectively and almost simultaneously, we adapted the stochastic synchronization computational framework of (Sarfati et al. 2023) which is based on the integrate-and-fire model of random oscillators. Our exploration of the model revealed that introducing a (fixed) refractory period after the breathing event is required to generate a high degree of synchronization in both homogenous and heterogeneous collectives of stochastic agents. This matches with the observation of the live shoal where individuals seem to be unresponsive for social cues of their neighbors once their old air is expelled and until they turn back into the deeper water with a c-startle (see SI Video 2). Here, individual fish tracking in 3D (Francisco et al. 2020) might help to pinpoint the exact time frame during which neighboring cues are ignored and thus future studies that apply such or similar techniques can help to add also spatial information of the individuals in the shoal to our understanding.

Our analysis further revealed that to match a Laplace-like distribution of the breathing intervals, the shoal must be composed of individuals with among-individual differences in own intrinsic breathing intervals as exemplified in simulated shoals with either heterogeneous or homogenous member compositions. The resulting peak of the breathing interval distribution in a heterogeneous group aligned with the most frequent interval of the fastest breathing type. This means that we observe a convergence of the collective breathing frequency to the intrinsic frequencies of the fastest breathers as those fast breathing individuals have shortest intervals between subsequent breathings and thus possess the narrowest window for a compromise in the breathing intervals. Fast breathers simply cannot stay longer underwater without breathing and the social coupling of their members results in many individuals following them to the surface. Those fast breathers may therefore be seen as key stone individuals (Modlmeier et al. 2014) with the strongest influence on the groups’ performances, while our simulations also show that their proportion in the group must be relatively small at about 10%. Increasing the proportion of fast breathers above 10% narrows the collective breathing distribution, reducing the “emergent variability” and making the group’s breathing pattern potentially easier to predict for predators. Our results are consistent with the previous work on leadership in groups (e.g., (Couzin et al. 2005)) where it was shown that a very small proportion of informed individuals (∼10%) is needed to guide the group decisions. Our numerical simulations also suggest that the real fastest-breathing individuals have not been included in our subset of individually tested fish, where we randomly picked 6 individuals from the shoal and from which we use the data to approximate breathing types in the simulations. While the model successfully replicated the Laplace-like shape of the empirical distribution, the location of the peak remained shifted compared to the empirical data (*µ* = 22.16s vs. *µ*_emp_ = 14.22s). Overall, our simulations suggest that shoals can only harbor a certain fraction of fast breathers in order to still possess a somewhat unpredictable surfacing frequency, and thus balancing selection on intermediate breathing types may come into play.

We observed that not all individuals took part in the collective air-breathing, although sometimes more than 80% or even almost all of the about 200 shoal members surfaced together. As a potential mechanism for this partial participation, we propose that individuals spatially self-assort based on their individual breathing needs (in our simulations implemented as fast, medium, slow breathing types). The non-random spatial clustering of individuals with similar behavioral traits has been previously studied in both animal and human groups (Aplin et al. 2013, Kurvers et al. 2014, Farine et al. 2015). Arapaima shoals are often layered resembling a moving cube or wall (“wall-like swimming mode”, see SI Video 1) where the upper layers are closest to the surface and - based on our qualitative observations - surface together while the bottom layers do not participate in the current collective breathing event. Such a self-organization might create spatial constraints, leading to stronger social coupling within the same breathing type (a cluster) and weaker social coupling between individuals that are spatially distinct and possibly possess a different breathing type. The implementation of this cluster interaction mechanism allowed us to reproduce part-taking distributions similar to the experimentally observed ones during the collective air-breathing. As the breathing intervals continue to follow Laplace-like distributions it can be assumed that this interaction mechanism modifies network structure dynamics while preserving the group-level patterns of behavior. Most interestingly, we found a clear transition region at relatively low inter-cluster coupling strength (*β*_*inter*_*clu*__ ∈ [0.6,1.0]) where every breathing type in the shoal can adjust partaking rates at almost the same compromise level. This means that in this region, even minute changes in responsiveness lead to immense changes in partaking rates while individual compromise is maintained at a similar level for all simulated breathing types. This aligns with principles of criticality, which propose that collective systems operate near critical points – parameter regime where small changes in one variable can lead to significant changes in system-wide behavior (Klamser and Romanczuk 2021, Poel et al. 2022, Gómez-Nava et al. 2023). In this regard, our model shows that by tuning the intra-cluster responsiveness between individuals of different behavioral types, a heterogeneous group can self-adjust to balance the inherent trade-offs between minimizing individual compromise and maximizing collective partaking rate. We believe that such mechanism can offer insights not only into biological coordination but also into the design of decentralized control in robotic swarms under diverse individual constraints.

We acknowledge the limited sample size of our empirical part of the study. However, as Arapaima in South America or Southeast Asia are bred in outside facilities with little to no possibilities to observe their behaviors underwater, our continuous four-hour recordings of a shoal represent a unique data set and thus a fruitful starting point for the investigation of this 23 Mio years old species. Based on the species’ old phylogenetic age, it is exemplified that collective behavior must have been evolved early on and similar assumptions have been drawn from fossils of highly polarized fish (Mizumoto et al. 2019). In future work, spatially explicit modelling combined with recording of the 3D positions of the individual fish in the tank would provide experimental evidence for the network structure dynamics during the socially coordinated air-breathing. Also, more systematic variation of live shoal compositions and shoal sizes would allow further verifying of the predictions derived from our simulations.

## Acknowledgement

The authors are grateful to Marcus Ebert, Laura Klatt, David Lewis, Mathias Kunow, and the “Team Technik” for their support at IGB. Furthermore, we thank Christopher Schutz, Nele Russy, Dustin Lehmann, Luis Gomez-Nava, Juliane Lukas, Olivia O’Connor, Manich Food Productions, Prof. Dr. Lenin Arias-Rodriguez, Matthias Stöck, and the Team of the Zoo Aquarium Berlin for their comments and support throughout the experimental process of this study. This research was partly funded by DFG-overheads for projects BI1828/2-1 and BI1828/3-1 (to DB). PB, JK, and DB acknowledge funding by the Deutsche Forschungsgemeinschaft (DFG, German Research Foundation) under Germany’s Excellence Strategy - EXC 2002/1 “Science of Intelligence” - project number 390523135.

## Supplementary Materials

**Fig. S1.**
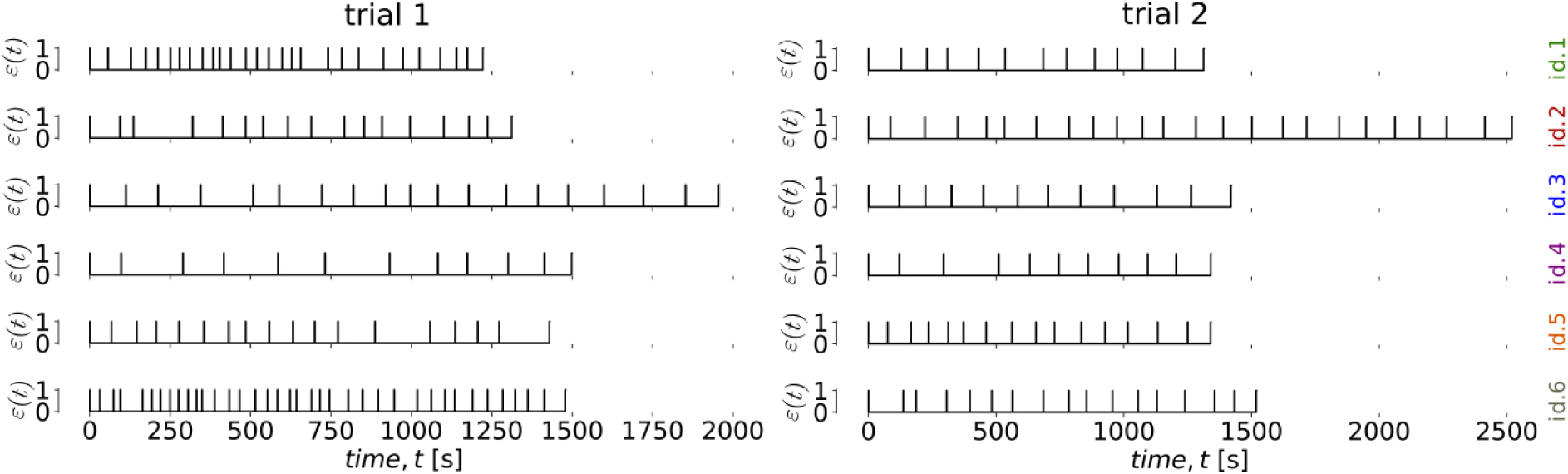
Time series of the individual breathing behavior of six individual fish recorded in isolation. Each spike, i.e., *ε*(*t*) = 1, depicts a breathing event of a single fish.

**Fig. S2.**
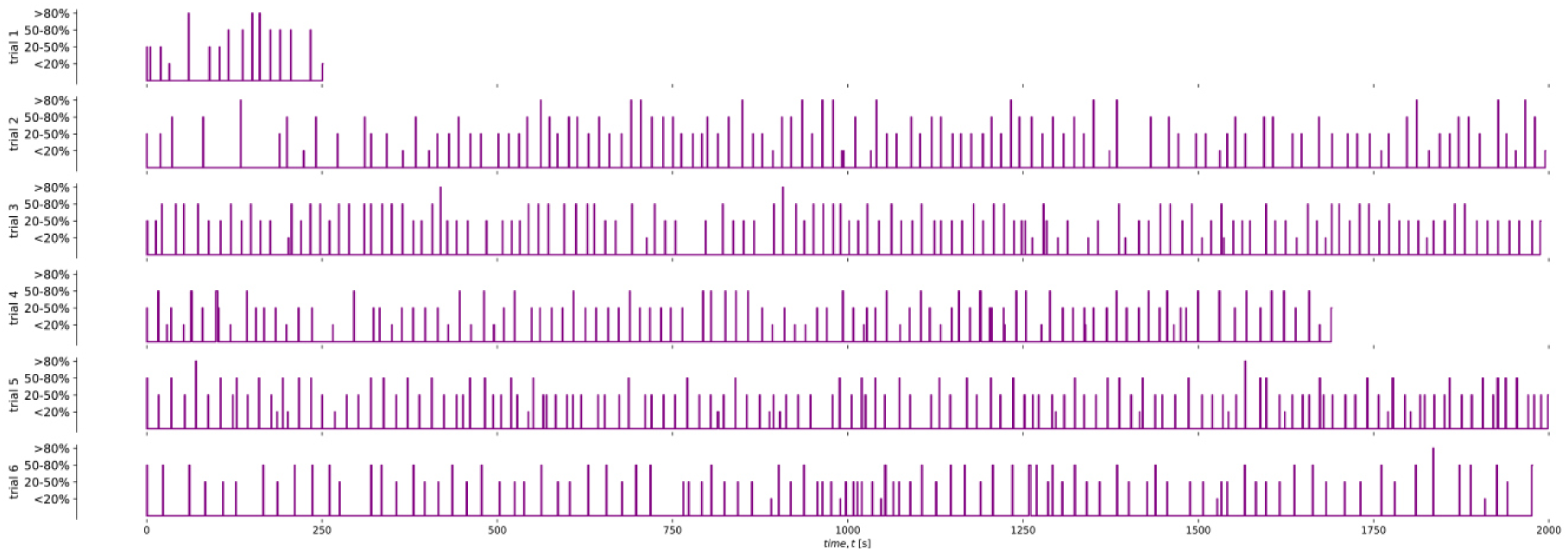
Time series of the collective breathing behavior of the investigated Arapaima shoal. Each spike depicts a collective breathing event, with a certain proportion of the shoal partaking (estimated as <20%, 20-50%, 50-80%, and >80% of the shoal).

**Fig. S3.**
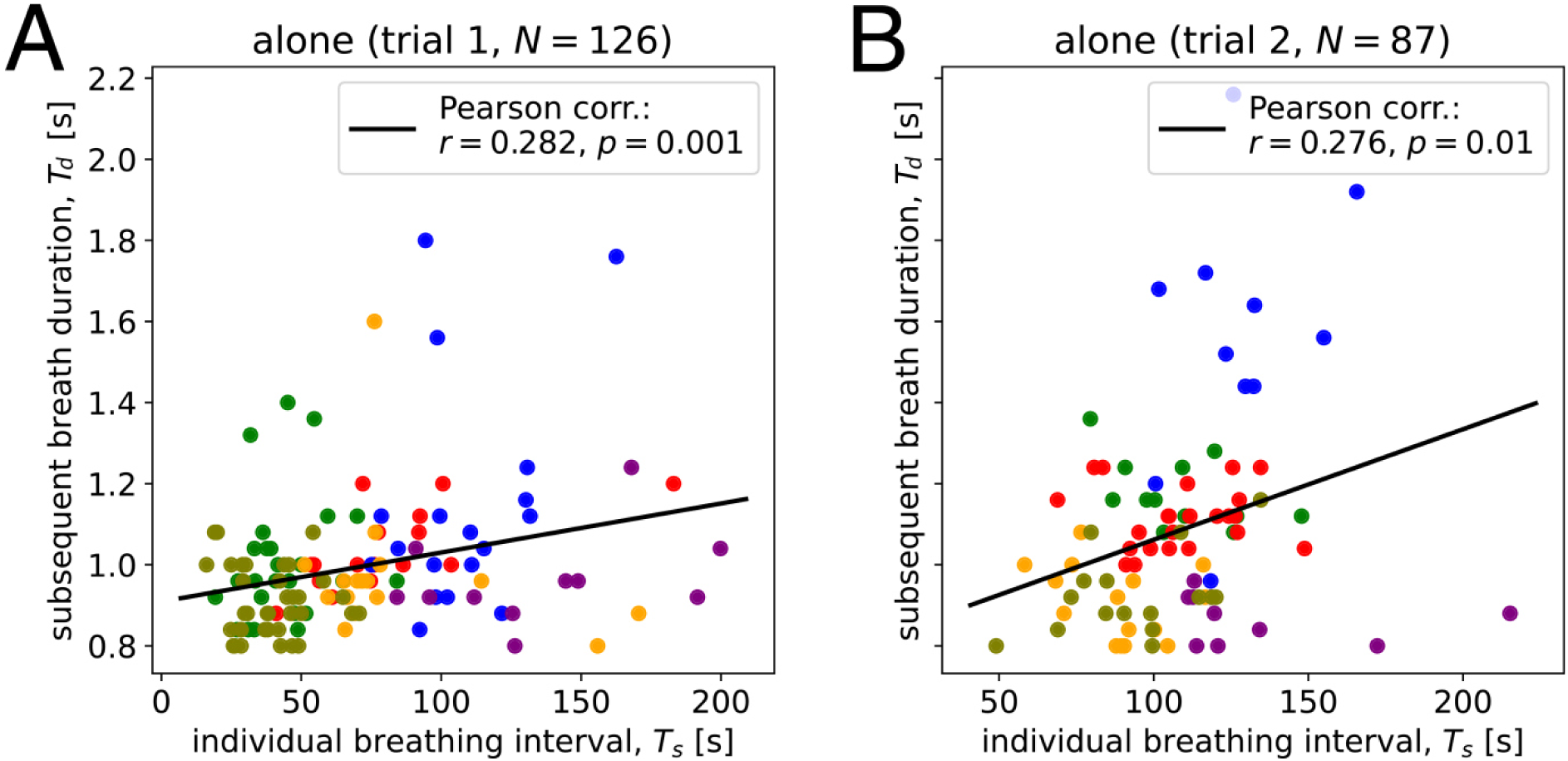
**Scatter plot of individual breathing intervals in isolation versus subsequent breath durations, with a fitted regression line for (A) trial 1 and (B) trial 2**. Dots of the same color correspond to the same individual. The positive correlation in both trials (trial 1: *r*=0.282, *P*<0.00; trial 2: *N*=87, *r*=0.276, *P*=0.01) suggests that a longer time since the last breath is associated with a longer subsequent breathing duration.

**Fig. S4.**
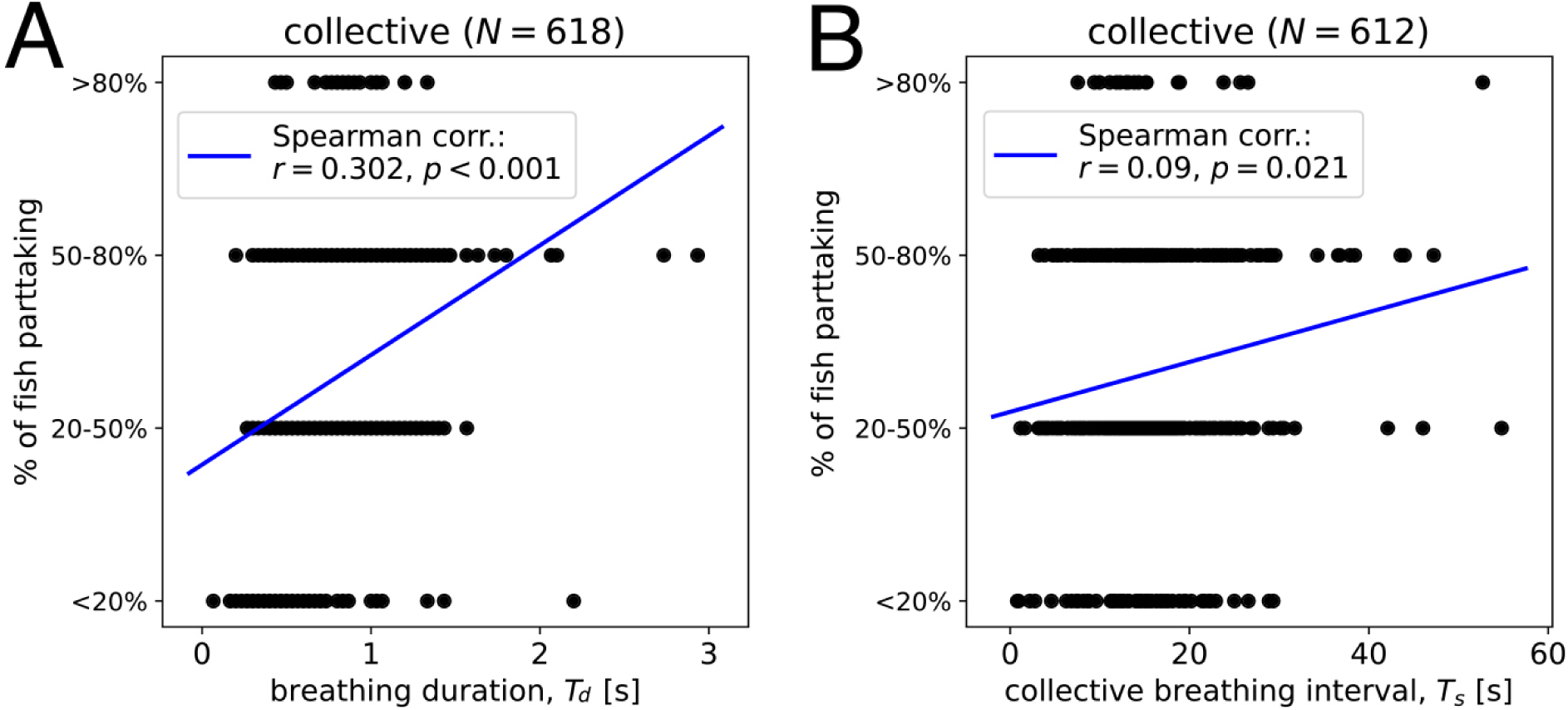
Scatter plots showing the relationship between the proportion of the shoal partaking in the breathing event and (A) the breathing duration, (B) the next collective breathing interval. The positive correlation (Spearman’s rank coefficient *r*_duration-partaking_ = 0.302, *p*<0.001) in (A) suggests that more fish taking part in the breathing event is associated with a longer breathing duration. The low Spearman’s rank coefficient (*r*_interval-partaking_ = 0.09, *p*=0.021) in (B) indicates that there is little to monotonic relationship between the proportion of the shoal partaking and the subsequent collective breathing interval.

**Fig. S5.**
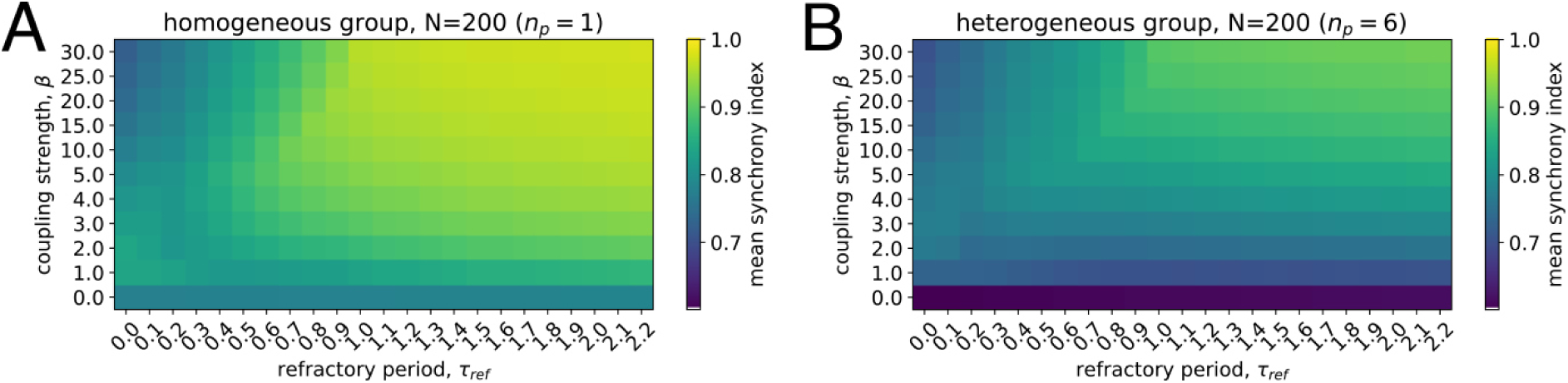
Synchronization in simulations. Mean synchrony index of breathing events depending on the strength of social interactions *β* and refractory period *τ*_*ref*_ in a homogenous (consisting of one breathing type with id=5) and heterogeneous (consisting of six breathing types) group of N=200 agents. The results were generated from 10 simulations, 1000 time steps each with *dt* = 0.01.

**Fig. S6.**
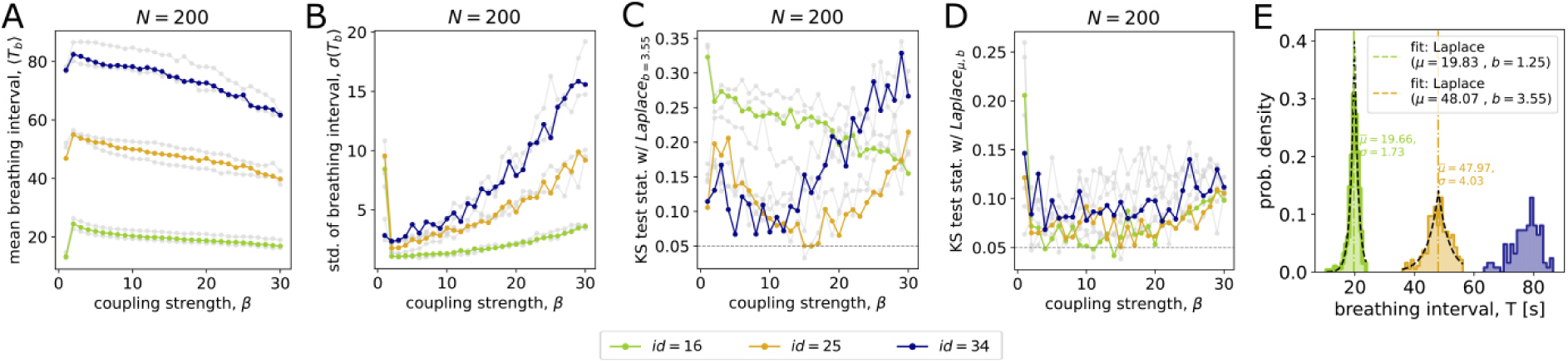
Simulated semi-heterogeneous groups, each composed of the individuals of two breathing types. (group id=16 in green: with 100 agents of id=1 and 100 agents of id=6; group id=25 in yellow: with 100 agents of id=2 and id=5; group id=34 in blue: with 100 agents of id=3 and 100 agents of id=4). (A) Mean and (B) standard deviation of the collective breathing intervals depending on the strength of social coupling *β*. Two-sided Kolmogorov-Smirnov test results verify at each β whether breathing intervals in the modelled shoal come from (C) the Laplace distribution with the empirical scale parameter *b* = 3.55 or from (D) any Laplace-like distribution without fixed parameters. (E) Depiction of the model-generated collective breathing interval distributions for three semi-heterogenous independent groups with β parameters corresponding to their best fits to the Laplace distribution (β=14 for a group with id=16 and β=15 for groups with ids ∈(25, 34)). The results in gray color in (A-D) correspond to homogeneous group compositions (for details see Fig. 4B-E in the main text) and are depicted for comparison.

**Fig. S7.**
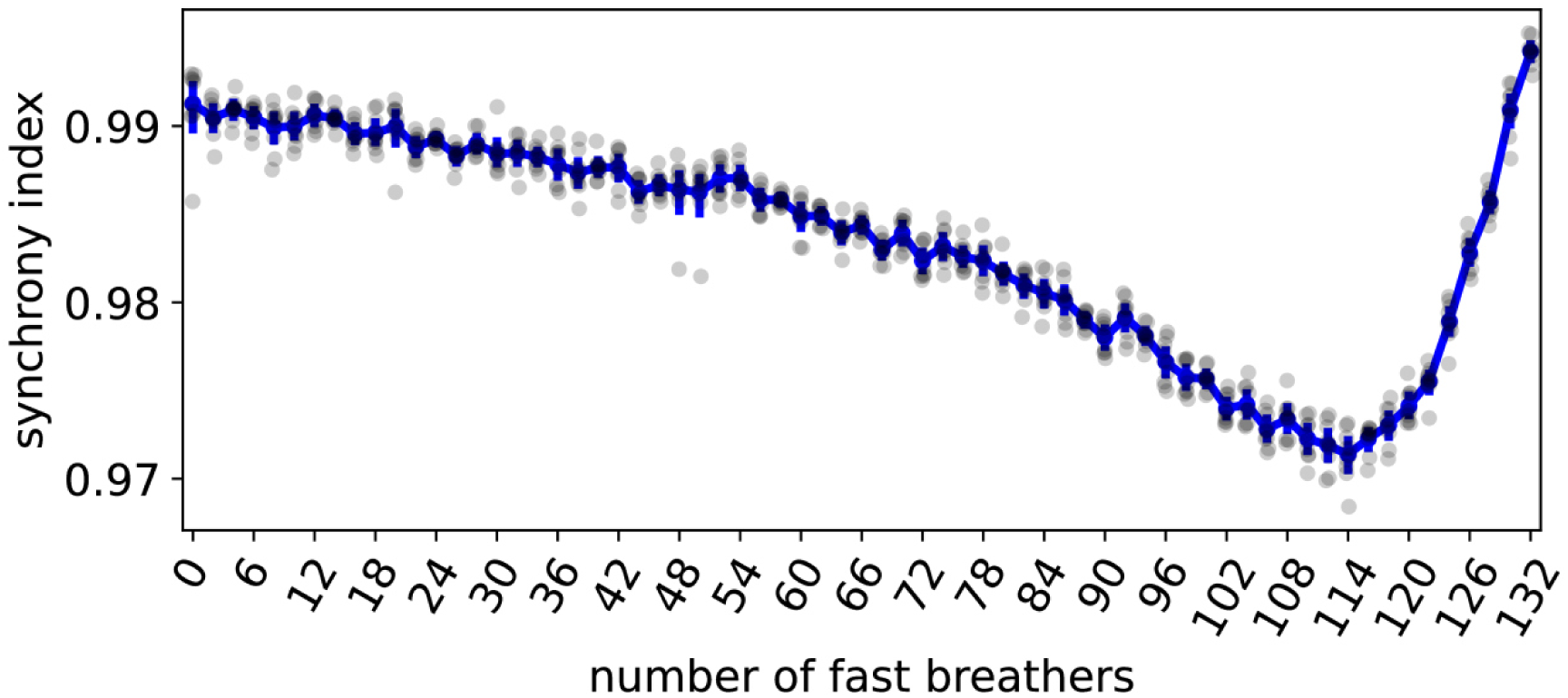
Impact of the number of fast breathers on the collective breathing synchrony in a simulated group of N=200. The number of present individuals of each of the breathing types (“fast”, “medium”, and “slow”) is re-calculated as (*N*_*fast*_, ⅓*N, ⅔*N − *N*_*fast*_), respectively. This way, the proportion of the medium breathers is kept constant to 34%, while the proportions of fast and slow breathers vary. The results were generated from 10 simulations, 1000 time steps each with *dt* = 0.01, social coupling strength of *β* = 14 and the refractory period *τ*_*ref*_ = 1.1. The mean synchrony index decreases as the number of fast breathers increases from 0 up to 114 (57% of the group, with 8% slow breathers) but increases afterwards, reaching full synchrony at 132 fast individuals (66% the group, with no slow breathers). Overall, synchrony of individual breathing events remains high within the range of 0.97 to 1.

**Fig. S8.**
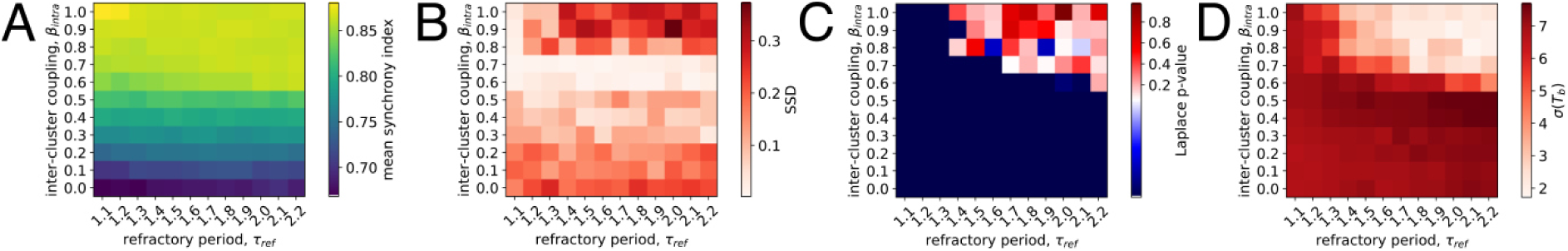
Impact of the refractory period on simulated collective breathing patterns’ characteristics depending on the inter-cluster coupling strength. *β*_*inter*_*clu*__ . (A) Mean synchrony index of breathing events across different combinations of refractory period and inter-cluster coupling strength. (B) The sum of squared differences (SSD) between empirical and simulation-generated probabilities of the proportions of individuals participating in breathing events. (C) Likelihood of encountering a Laplace-like distribution of collective breathing intervals depending on the refractory period and inter-cluster coupling strength. If the p-value is below 0.05 (in blue), the intervals do not follow a Laplace-like distribution. (D) The standard deviation of the simulated collective breathing intervals for different combinations of refractory period and inter-cluster coupling strength. The results were obtained using simulations with heterogeneous three-personality groups of *N* = 200 individuals with ids ∈ (4,5,6) from 10 realizations, 1000 time steps each with *dt* = 0.01 and fixed intra-cluster coupling strength of *β* = 15. Based on the results in the main text, we assumed fast breathers are in the minority and represent the group with an unequal ratio of fast:medium:slow breathers as 1:2:2, corresponding to cluster sizes of *N*_*clu*_ ∈ (40,80,80) agents.

**Table S1.** Proportion of simulated breathing events where individuals of certain type (i.e., “slow”, “medium”, “fast” breather) were breathing alone without synchronizing with a collective. The results are reported as a function of inter-cluster coupling strength *β*_*inter*_*clu*__ for different breathing types. The proportion is computed relative to the total number of breathing events from 10 simulations, 1000 time steps each with *dt* = 0.01, intra-cluster coupling strength of *β*_*intra*_*clu*__ = 15 and refractory period *τ*_*ref*_ = 2.1. The values correspond to the fraction of data points in Fig. 6 of the main text where the breathing group ratio equals 0 (i.e., the individual took breath alone) relative to the total number of points (breathing events) for the corresponding *β*_*intra*_*clu*__. The values in red are in the transition region, where the collective breathing interval distribution best matches the Laplace-like distribution and empirical shoal partaking rates.

| $\beta_{inter_{clu}}$ | slow breather | medium breather | fast breather |
| --- | --- | --- | --- |
| 0.0 | - | 0.01 | 0.05 |
| 0.1 | 0.06 | 0.01 | 0.06 |
| 0.2 | 0.03 | 0.02 | 0.07 |
| 0.3 | 0.05 | 0.01 | 0.11 |
| 0.4 | 0.07 | 0.01 | 0.12 |
| 0.5 | 0.09 | 0.09 | 0.13 |
| 0.6 | 0.01 | 0.09 | 0.32 |
| 0.7 | - | 0.01 | 0.17 |
| 0.8 | - | 0.01 | 0.04 |
| 0.9 | - | - | 0.01 |
| 1.0 | - | - | 0.004 |
| 1.5 | - | - | - |
| 2.0 | - | - | 0.03 |
| 2.2 | - | - | 0.12 |
| 2.5 | - | - | 0.40 |
| 2.8 | - | - | 0.57 |
| 3.0 | - | - | 0.32 |
| 4.0 | - | - | 0.01 |
| 5.0 | - | - | 0.002 |
| 15.0 | - | - | 0.002 |

**Supplementary note 1: Agent-based simulations with homogenous group compositions.**

Using computer simulations, we found that collective breathing intervals in homogeneous groups did not fit a Laplace distribution with the empirical scale parameter b, except for a group composition with individuals of id=2 (see in the main text, Fig. 4D). While homogeneous groups of id=2 with high coupling strength *β* = 15resembled the empirical collective breathing interval distribution in shape, a high mean and location parameter (*µ* = 44.79) indicated a significant deviation from the experimental data (see in the main text, Fig. 4А, F). Homogeneous groups of id=1 or id=6 were both described by smaller mean of breathing intervals compared to others, but did not fit the Laplace distribution of collective breathing intervals with the empirical scale parameter (see in the main text Fig. 4A, D). For instance, homogenous groups of id=1 generated Laplace-like distributions of collective breathing intervals with location parameter of *µ* = 22.18 but had a narrower shape (*b* = 1.11) than the experimental distribution (see in the main text Fig. 4E, F). Meanwhile, the breathing patterns of other homogeneous group compositions did not reproduce a Laplace-like distribution of collective breathing intervals in general (Fig. 4E).

**Supplementary note 2: Agent-based simulations with semi-heterogeneous group compositions.**

As a special case of heterogenous group composition, we analyzed the collective breathing dynamics of semi-heterogeneous groups. Each semi-heterogenous group (N=200) consisted of an equal proportion of individuals from two breathing types which characterized by close mean breathing intervals: group (id=16) with 100 individuals of id=1 and 100 individuals of id=6; group (id=25) with 100 individuals of id=2 and 100 individuals of id=5; group (id=34) with 100 individuals of id=3 and 100 individuals of id=4. We found that semi-heterogeneous compositions closely resemble the collective dynamics of their homogeneous counterparts (see Fig. S6A-C). This suggests that the inter-individual breathing differences between the constituting types within each composition (although statistically consistently differ for single individuals in isolation) are not strong enough to significantly impact group behaviour. This may also imply that, while breathing types vary on a continuous scale, the most impact on collective breathing dynamics comes from “category-like” differences between individuals. Overall, the results showed that the mean breathing interval in semi-heterogeneous groups was shifted towards the faster breathing type (Fig. S6A, Fig. S6E vs. Fig. 4F in the main text).

Notably, when a breathing type that followed a Laplace distribution of the breathing intervals in a homogeneous shoal (e.g., id = 1 or id =2, see Fig. 4E-F in the main text) was mixed with another breathing type that did not follow a Laplace distribution in a homogeneous shoal, the collective breathing interval distribution of such a two breathing types mixed group remained Laplace (see e.g., id=25 or id=16 in Fig. S6D-E). This suggests that a subgroup (N=100) of a breathing type that followed a Laplace distribution of breathing intervals in a homogeneous case remains robust and even dominates, i.e., shapes the overall group breathing dynamics, when mixed with a close, yet distinct, breathing type. Indeed, a semi-heterogeneous group (id=25) with individuals of id=2 and id=5 followed a shape of empirical Laplace distribution (see Fig. S6C in orange, coupling strength *β* = 15), similarly to a homogenous group consisting of only individuals of id=2 (see in the main text Fig. 4D in red). Other semi-heterogeneous group compositions (id=16 and id=34) did not fit the Laplace distribution with the empirical scale parameter *b* (Fig. S6C), similarly to their homogeneous counterparts (see in the main text Fig. 4D).

**Supplementary note 3: Robustness analysis of simulated group synchrony.**

To assess the degree of synchronization of individuals’ breathing events in a simulated group (i.e., synchrony index), we used the spike_sync() function from the Python library “PySpike”, which measures similarity between individuals’ spike trains (Kreuz at al. 2015, Mulansky and Kreuz 2016). In out case, a spike train represents an array of the starting times of an individual’s ibreathing events (i.e., when *ε*_*i*_(*t*) = 1) over the duration of a single simulation (i.e., 10000 time steps).

We tested the robustness of our agent-based model against the refractory period *τ*_*ref*_ and coupling strength *β* in homogenous and heterogeneous groups (see Fig. S5). We found a transition from lower to higher synchrony with increasing coupling strength and refractory period in both group compositions. Both groups exhibited a high degree of synchronization when the refractory period was non-zero. Notably, for *τ*_*ref*_ > 1.0, the degree of synchronization remained robust with no further changes, regardless of whether the group was homogenous or heterogeneous. Based on empirical observations of high synchronization in collective breathing events, we set *τ*_*ref*_ = 1.1 in our simulations.

We also tested how tan unequal distribution of breathing types affects the degree of synchrony in heterogeneous group compositions (Fig. S7). As the proportion of fast breathers increased (up to 57%) and slow breathers decreased (down to 8%), overall synchrony gradually declined from 0.99 to 0.97. Interestingly, when slow breathers were further reduced to 0%, the group regained full synchronization. Overall, the synchrony of individual breathing events in a group remained high (i.e., within the range of 0.97 to 1), suggesting robustness to group composition imbalances in the distribution of breathing types.

In a scenario with cluster synchronization (see Fig. S8), where only fractions of agents rather than the entire group participated in the collective breathing events, we tested the robustness against the refractory period *τ*_*ref*_ and the inter-cluster coupling strength *β*_*inter*_*clu*__. We found that the degree of synchronization increased with higher inter-cluster coupling strength and remained robust across the tested range of the refractory period (Fig. S8A).

The sum of squared differences (SSD) between empirical and simulation-generated probabilities of the proportions of individuals participating in breathing events was lowest in the transition region *β*_*inter*_*clu*__ ∈ {0.6,0.7}, and maintained robustness across different values of the refractory period (Fig. S8B). Interestingly, both the likelihood of encountering a Laplace-like distribution of collective breathing intervals and their standard deviation were sensitive to changes in the refractory period (Fig. S8C,D). Therefore, to match the empirical observations both in terms of SSD and the emergence of Laplace-like distribution of collective breathing intervals, we set the refractory period to *τ*_*ref*_ = 2.1 in our partaking simulations.

